# Characterization of convergent thickening, a major convergence force producing morphogenic movement in amphibians

**DOI:** 10.1101/270892

**Authors:** David Shook, Jason Wen, Ana Rolo, Brian Francica, Destiny Dobins, Paul Skoglund, Doug DeSimone, Rudolf Winklbauer, Ray Keller

**Affiliations:** Department of Biology University of Virginia, Charlottesville, VA 22904; Department of Cell and Systems Biology, University of Toronto, Toronto, Ontario Canada M5S 3G5CA; Centre for Craniofacial and Regenerative Biology King’s College London London SE1 9RT, UK; Aduro Biotech 740 Heinz Ave. Berkeley, CA 94710; 52826 Cedar Street Philadelphia, PA, 19134; Department of Cell Biology University of Virginia, School of Medicine Charlottesville, VA 22908

## Abstract

We characterize the morphogenic process of convergent thickening (CT), which occurs in the involuting marginal zone (IMZ) during gastrulation of *Xenopus*, the African clawed frog. CT was described previously as the tendency of explants of the ventral IMZ of *Xenopus* to converge their circumblastoporal dimension and thicken their radial dimension (Keller and Danilchik 1988). Here we show that CT occurs from the onset of gastrulation, initially throughout the pre-involution IMZ. We suggest that CT is driven by an increase in the interfacial tension between the deep IMZ and its epithelium, resulting in cells of the deep IMZ tending to minimize their surface area. In explants, this results in a progressive shortening (convergence) of the IMZ along its longer mediolateral axis and thickening in the orthogonal planes, and can generate tensile force (Shook et al. 2018). In vivo, convergence of the annular IMZ generates circumferential tension, closing the blastopore. These results provide the first clear example of a tensile morphogenic force from a Holtfreterian/Steinbergian change in tissue affinity.

## INTRODUCTION

Amphibian gastrulation involves the movement of the cells that will give rise to the internal tissues of the embryo from the surface to the inside of the embryo, thereby establishing the fundamental structure of the three-layered adult body plan. These movements depend on forces generated by these same cells within the embryo. One set of movements involves the “convergence” or decrease in circumference of the ring of cells comprising the involuting marginal zone (IMZ), which lies around the vegetal endoderm (VE) at the onset of gastrulation (Figure 1A, B), a convergence that generates a constricting force around the VE, closes the blastopore (Figure 1A), and ultimately helps to push both the VE and the IMZ inside the embryo.

**Figure 1.** The behavior and role of the involuting marginal zone (IMZ) in blastopore closure. The vegetal and animal limits of the IMZ are indicated by yellow and green lines superimposed on to vegetal views of a Xenopus gastrula stage embryo at the indicated Nieuwkoop and Faber stages (REF) (A). Formation of the blastoporal pigment line, which defines the vegetal end of the IMZ, is indicated by yellow pointers, and formation of a depression, the initiation of involution (and an invagination) of the IMZ is indicated by green pointers. Note that the vegetal boundary of the IMZ is carried inside (dashed yellow lines) and covered over by the blastoporal lips from dorsal to ventral as gastrulation proceeds. Explants of the dorsal quarter of the IMZ (B-C), including tissues of presumptive notochordal (magenta), somitic (red), and neural (blue) fates undergo convergent extension (CE, green-blue symbols, C), whereas explants of the ventral quarter (B-D), which develop only ventrolateral mesoderm (orange) when separated from dorsal tissues, round up and show a thickening of the mesodermal tissues, a movement called Convergent Thickening (CT, white symbols). Together, CE on the dorsal side and CT on the ventral side of the IMZ were previously thought to provide the convergence forces that close the blastopore (as shown in A) (Keller and Danilchik 1988). Here, a more detailed analysis of standard “giant” explants of the entire marginal zone (B-E), and variants thereof, reveals a revised view of the pattern of CT expression, wherein initially the entire IMZ uniformly expresses CT during early gastrulation (white symbols, E) and at the midgastrula stage expression of CT begins to transition into a progressive expression of CE (E’, E”), a postinvolution progression from anterior to posterior, as described previously (Shih and Keller 1992b), while CT persists in the preinvolution IMZ (E’, E”). Finally, ventralized embryos, which lack the presumptive tissues of notochordal and somitic mesoderm and contain only presumptive ventrolateral tissues, closed their blastopores symmetrically (green arrows, F), using CT alone (white symbols, F), and a giant explant of the ventralized embryo, consisting of entirely ventral tissues undergoes a uniform CT (G’). For a complete description of other types of explants, see Methods and Figure 1-supplementary figure 1.

Here we characterize the expression, the mechanism and the function of one of these movements, convergent thickening (CT), and define its relationship to the other convergence movement, convergent extension (CE) during gastrulation of the African clawed frog, Xenopus laevis. The blastopore forms with the appearance of a pigment line (yellow arrow heads, Figure 1A, stage 10) on the dorsal side at the lower boundary of the IMZ with the VE (yellow line, Figure 1A, stage 10). This pigment line forms by apical constriction, resulting in concentration of pigment in the apices of the IMZ cells bounding the VE. This apical constriction contributes to formation of the wedge shaped “bottle cells”, which bend the IMZ inward to form an invagination, (see Keller 1981; Hardin and Keller 1988). This sequence of events begins dorsally and progresses laterally and ventrally around the VE (yellow arrow heads, Figure 1A, stage 10-10.5) and is followed by initiation of involution (green arrow heads, stage 10.25-11, Figure 1A,).

During this process, the IMZ itself begins to converge around the blastopore. As it converges, the IMZ also moves through the blastopore by “involution”, or rolling over the lip and passing into the interior; the approach of the cells within the IMZ toward the blastopore can be tracked on the embryo (green line at stage 10 moves toward blastopore through stage 11, Figure 1A,). Note that by convention, the initial “blastopore” is defined by the blastoporal pigment line separating the IMZ from the VE. However, once the lips of involuting IMZ are formed, they, rather than the locus of bottle cell formation, defines the blastopore and blastopore closure is defined by the centripetal progress of the inner edge or “lip” of the IMZ across the VE as the pigment line forming the original boundary is covered over (dashed yellow line, stage 10.5-11, Figure 1).

A number of morphogenic mechanisms facilitate or contribute to the convergence of the IMZ and blastopore closure. First, vegetal rotation (VR), a morphogenic movement that involves the active, autonomous upwelling of the central VE and contraction of its exterior surface (Winklbauer and Schurfeld 1999; Ibrahim and Winklbauer 2001; Wen and Winklbauer 2017), which reduces resistance to the convergence of the IMZ, positions more of the VE higher within the sphere of the embryo and with respect to the converging ring of the IMZ, such that the converging IMZ tends to push the VE further inward rather than outward (exogastrulation). Second, epiboly, a morphogenic movement that involves the passive thinning and spreading of the animal cap region (Keller 1980), also reduces resistance to IMZ convergence; failure of epiboly can block blastopore closure (e.g. Rozario et al. 2009; Eagleson et al. 2015), whereas removal of the animal cap region can accelerate it (Keller and Jansa 1992). Third, the apical constriction of bottle cells at the vegetal edge of the annular IMZ tends to decrease the IMZ’s circumference somewhat (see Hardin and Keller 1988). Finally, once involution begins, the bottle cells are rapidly internalized, removing them from the context in which they can affect blastopore closure.

In addition, there are two intrinsic convergence behaviors of the IMZ itself that drive blastopore closure, and the best characterized thus far has been CE (Keller et al. 2000; Keller et al. 2008). CE is defined as an autonomous, force-producing morphogenic movement that narrows and lengthens the presumptive notochordal and somitic mesoderm of the dorsal sector of the gastrula in vivo (Vogt 1929; Keller 1976) and when explanted into culture (Schechtman 1942; Keller and Danilchik 1988) (Figure 1B-C), where the explanted tissues mimic the movements of those in vivo. The second convergent morphogenic movement, CT, was initially described as a tendency for explants of the ventral sector of the early gastrula to converge and thicken and form a mini-gastroid lacking dorsal tissues (Figure1B-D) (see Keller and Danilchik 1988). CE was previously thought to be the dominant convergence process driving blastopore closure, originating in the dorsal sector of the IMZ and operating exclusively in presumptive dorsal tissues (the presumptive notochordal, somitic, and posterior neural tissue), while CT was thought to operate only in the ventral IMZ (Keller and Danilchik 1988).

However, several long-standing observations suggest that this is not true. First, the highly anisotropic nature and the timing of CE makes it difficult to account for the largely symmetric convergence movements of the preinvolution IMZ during gastrulation (Keller and Danilchik 1988) and does nothing to explain the convergence around the entire blastopore during early gastrulation. CE of the mesoderm is a *post-involution* process of convergence driving lengthening of presumptive notochordal and somitic mesoderm, and begins two hours after the onset of gastrulation, initially operating only with the dorsal quadrant, and only reaching the tissues initially laying most ventrally after blastopore closure (see Shih and Keller 1992b; Shih and Keller 1992a; Lane and Keller 1997). Coupled with the coordinate CE of the overlying posterior neural plate (Keller 1978; Keller et al. 1991; Keller and Winklbauer 1992), mesodermal CE results in an asymmetric, dorsal-dominated eccentric closure of the blastopore of normal embryos toward the ventral side of the blastopore. However, excepting bottle cell formation, the *pre-involution* convergence movements of the IMZ are surprisingly isotropic all around the blastopore of the early and mid gastrula (Keller and Danilchik 1988). This then suggests that there may be an *isotropic* convergence process operating within the preinvolution IMZ throughout gastrulation. Second, Xenopus embryos ventralized by UV irradiation develop only ventral tissues, and thus lack the dorsal tissues (notochordal and somitic mesodermal and posterior neural tissues) that express CE. Nevertheless, they close their blastopores (first shown by Scharf and Gerhart 1980) and do so symmetrically, showing consistently isotropic convergence of the IMZ (our results), suggesting that they use CT to close the blastopore without CE. Third, the fact that normal embryos of other amphibians, such as Gastrotheca, close their blastopore symmetrically without expression of dorsal tissue markers and delay CE until after blastopore closure (del Pino and Elinson 1983; del Pino 1996; Benitez and Del Pino 2002) suggest that CT is also responsible for BP closure in these species, and that CT may be a common feature of amphibian gastrulation whereas CE is used with different timings in different species (Keller and Shook 2004; del Pino et al. 2007).

Here we characterize the pattern of tissue deformation, the spatiotemporal pattern of expression, the regulation, and the cellular mechanism of CT in normal and ventralized Xenopus embryos. We show that: 1) CT is expressed from the onset of gastrulation, before CE; 2) it is expressed not only in the ventral sector of the IMZ but circumferentially, in all sectors of the IMZ of normal embryos (Figure 1E); 3) that it occurrs outside the blastopore, in the portion of the IMZ that has not yet involuted; 4) tissues expressing CT undergo transition to expressing CE in explants (Figure 1E-E”), a transition that occurs at involution in the embryo. Our early experiments trying to discern the cell behavior underlying CT revealed no polarized cell behavior. Because explanted IMZ tissue tends to round up and shows a loss of affinity between its superficial epithelial layer and the underlying deep, mesenchymal, mesodermal tissues at the onset of gastrulation, we hypothesized that tissue surface tension (Steinberg and Poole 1981; Forgacs et al. 1998; Lecuit and Lenne 2007) could be driving some of the observed CT in explants, while this tendency generates circumferential tension in intact embryos. We further show that: 5) whereas the interfacial surface tension between the deep and epithelial layers of ectodermal tissues stays constant, surface tension between the deep and epithelial layers of the IMZ increases at gastrulation, consistent with its playing a role in thickening and convergence of the deep tissue; 6) CT is expressed throughout gastrulation in the IMZ of ventralized embryos (Figure 1F-G’) and is the major force driving their symmetric blastopore closure. Elsewhere we show that CT of the IMZ is capable of generating early convergence forces during normal gastrulation, prior to the onset of CE, and that IMZs of ventralized embryos can generate convergence forces approximating those of the combination of CE and CT in normal embryos (Shook et al. 2018). Taken together, these results support the hypothesis that CT and CE are two widely expressed, sequential convergence mechanisms contributing to blastopore closure and gastrulation in Xenopus laevis. The significance of these results for the use of tissue surface tension-based mechanisms in morphogenesis and for the evolution of gastrulation and the likely significance of CT in other organisms are discussed.

## RESULTS

### Characterization of the early expression of CT in normal and ventralized embryos

Pre-involution convergence of the IMZ of normal embryos occurs at a similar rate in the dorsal, lateral and ventral regions around the blastopore through the first two hours after the onset of gastrulation at stage 10, although it begins earlier dorsally (Figure 2B), associated with the onset of bottle cell formation there an hour before it begins to progress around the lateral and ventral side of the blastopore (2A, left column, stage 10-10.5; Movie A).

**Figure 2.**
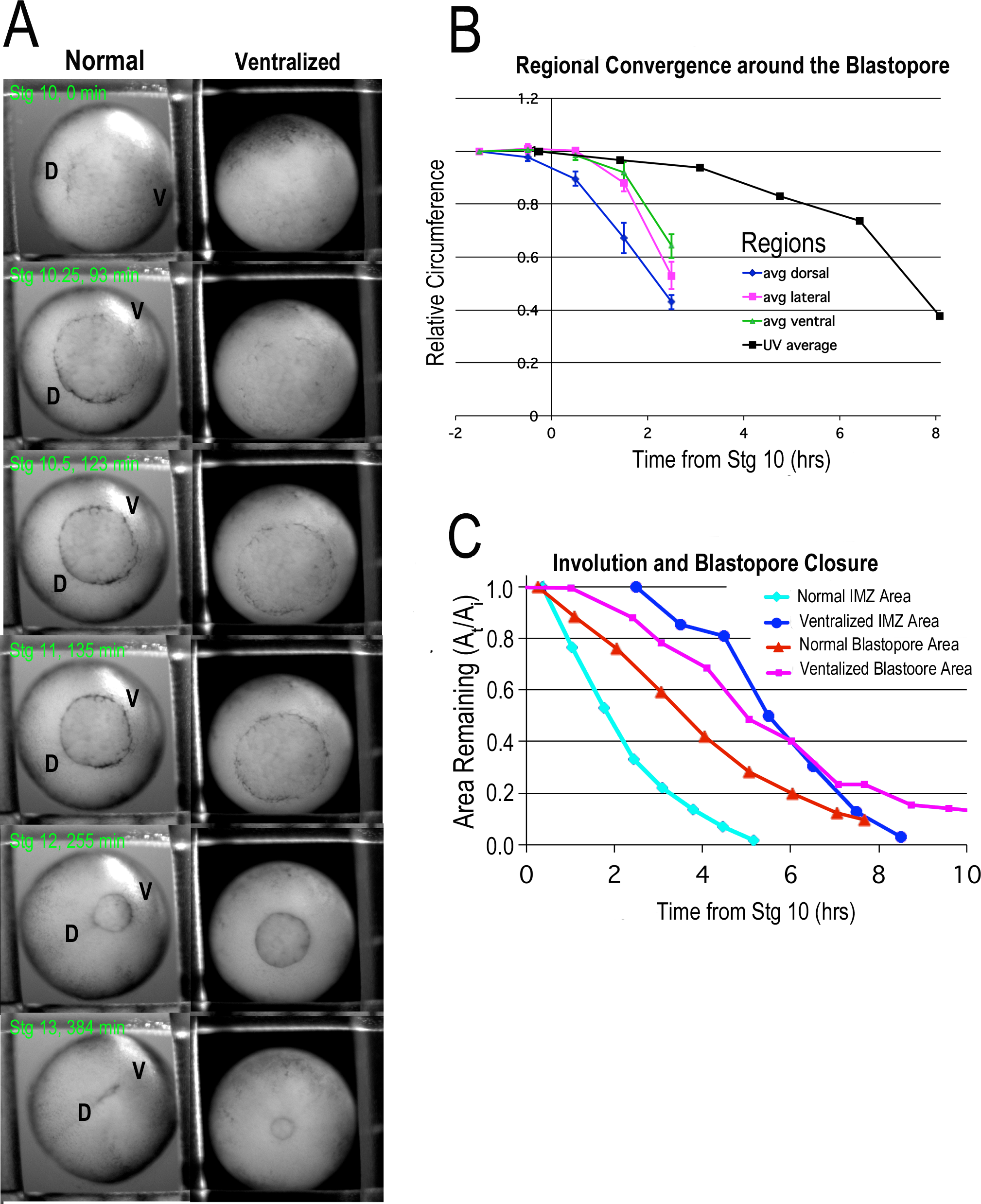
Time lapse movie frames of normal (left) and ventralized embryos (right) are shown at the stages and times indicated (A). Note the blastopore formation (blastopore pigment line formation) and subsequent blastopore closure is delayed in the ventralized embryos, with formation of the blastopore usually occurring at control stage 10.5. Note also that the blastopore of ventralized embryos closes symmetrically whereas the normal closure is biased toward the future ventral side (“V”, upper right in A). Analysis of rates of convergence in the dorsal, lateral and ventral quadrants of the preinvolution IMZ of normal embryos shows similar rates of convergence, with an earlier onset in the dorsal quadrant (B). Analysis of change in superficial area of the pre-involution IMZ compared to its original size, a measure of its involution, and change in area of the blastopore (the area of the exposed vegetal endoderm), a measure of blastopore closure, from timelapse tracings (C).

It is only from the mid-gastrula onward that blastopore closure shows the anisotropic effects of mechanical coupling to the postinvolution mesodermal and posterior neural tissues undergoing the highly anisotropic CE, which generates greater extension dorsally (see Discussion and Keller and Danilchik 1988) and thus biases the closure of the blastopore toward the ventral side (Figure 2A, left column, compare stage 10.5 to 12; Movie A).

Embryos ventralized by UV irradiation of the vegetal hemisphere before first cleavage (see Scharf and Gerhart 1980) begin to form bottle cells nearly simultaneously all around the embryo at about control stage 10.5 (Figure 2A, right column, stage 10,5), a time when normal embryos form bottle cells ventrally (Figure 2A, left column, stage 10.5). Convergence occurs uniformly around the IMZ, and blastopore closure occurs symmetrically around the VE (Figure 2A, right column; Movie A), but begins later and proceeds more slowly (Figure 2B).

Rates of blastopore closure resulting from convergence of the IMZ, based on the area of the exposed VE, are similar in normal and ventralized embryos, but show a delay of 1-2 hours in the extent of BP closure in ventralized embryos (Figure 2C), consistent with the delay in bottle cell formation and in convergence seen in ventralized (Figure 2A,B). Involution of the IMZ in ventralized embryos, on the other hand, begins more slowly compared to normal embryos and is delayed by 3-4 hours (Figure 2C). This disparity in convergence and VE internalization vs. involution (mesoderm internalization) may be because VR cooperates with CT to internalize the VE, while CT is solely responsible for involution in ventralized embryos. The implication would be that CT can replace CE to generate circumblastoporal force (Shook et al. 2018) to drive BP closure, but is not as effective as CE at driving involution. The slow involution of ventralized embryos leaves a portion of the IMZ in the un-involuted position for far longer. Assuming this also delays a transition from CT to another cell behavior, this would allow CT to generate circumblastoporal force for longer. This is also likely the case in anuran species that delay CE until late gastrulation or neurulation. The opposite result is true for many urodele species, where involution occurs much more rapidly than BP closure.

These results demonstrate that convergence and blastopore closure begins well before the onset of CE, around the entire circumference of the IMZ, and that CT can help drive BP closure, as well as involution, albeit more slowly on its own. It also explains the largely isotropic convergence of the pre-involution IMZ around the circumference of the BP.

### CT is autonomous, begins prior to CE and acts throughout the entire preinvolution IMZ

Explanting tissues from the embryo allows us to isolate morphogenic movements from their geometric constraints within the intact embryo and the influence of other tissues, and to thereby determine which mechanical deformations result from autonomous processes of the tissue itself. In addition, explantation makes the tissues more accessible to imaging and measurement. To test the autonomy and dynamics of CT, we isolated the entire non-involuting marginal zone (NIMZ) plus IMZ from stage 9 embryos as giant sandwich explants (Figure 1E), isolated with or without the presumptive bottle cells (Figure 3A, B; Movies D, F; Figure 1-supplementary figure 1A,B). Both types of explants show strong convergence of the IMZ (lower, less densely pigmented region), indicating that the bottle cells play, at most, a minor role in early convergence movements of unrestrained explants. The NIMZ consistently converges to a lesser extent than the IMZ after stage 10- (data not shown), such that the explants consistently bow toward their vegetal (bottom) end. To determine whether there was any difference in the amount of convergence dorsally vs. ventrally, suggestive of a stronger regional expression of CT, or in the onset of convergence, suggestive of a dorsal to ventral progression of CT, we plotted regional convergence of the widest portion of the IMZ across its mediolateral extent in standard, dorsally-centered giant explants (Figure 3C) and also, as a control for healing artifacts at the lateral edges, in ventrally-centered giant explants (Figure 3D; see Methods, Figure 1-supplementary figure 1C). In both cases, convergence in dorsal, lateral and ventral regions (examples of these regions indicated by arrowheads, Figure 3A) show no differences between sectors in the first two hours of gastrulation, prior to the onset of CE (see also Movies D, W), despite differing times of bottle cell formation. Convergence rates in dorsally and ventrally centered giant explants were similar to those observed in giants made from ventralized embryos (Figure 3E, Figure 1-supplementary figure 1F), which show continuous CT of the IMZ for several hours after explantation (Figure 3-supplementary figure 1A; Movie E). Giant sandwich explants made from Li+ dorsoanteriorized embryos show isotropic convergence both before and after they begin to extend (Figure 3-supp. figure 1B; Movie G; Figure 1-supplementary figure 1G).

**Figure 3.**
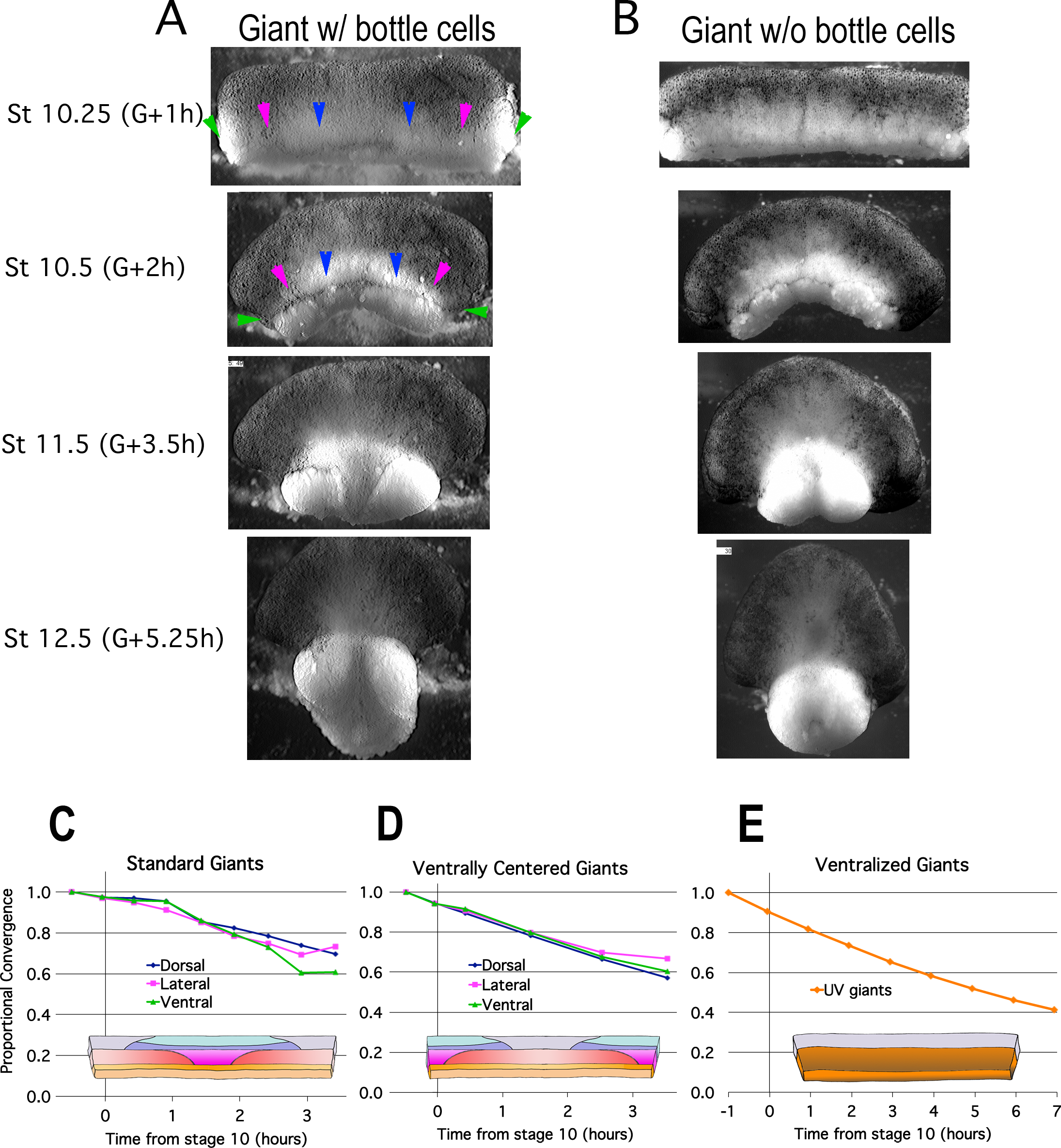
Lack of dependence on bottle cells & uniformity of regional convergence. Frames from time lapse movies of giant sandwich explants with (A) and without bottle cells (B) after release from their constraining coverslips show uniform convergence of the involuting marginal zone, with no dependence on bottle cells to drive convergence. To compare regional rates of convergence in such explants, the mediolateral extent of the dorsal (between blue pointers), lateral (between magenta and blue pointers), andventral regions (between green and magenta pointers) (A) was measured at time points from stage 10− and plotted with respect to initial width (W_t_/W_i_) in standard (dorsally centered) giants (C), ventrally centered giants (D) and giants from ventralized embryos(E).

These results indicate that CT is an intrinsic tendency of the entire IMZ prior to the onset of CE and shows no bias in timing or extent with regard to dorsal-ventral patterning, and that early convergence is not dependent on bottle cell formation. CT is expressed in the pre-involution IMZ, regardless of whether CE is later expressed in its normal pattern (in the presumptive anterior to posterior axis, which runs from the dorsolateral to the ventrolateral sectors of the gastrula), is not expressed at all in ventralized embryos, or is later expressed everywhere at the same time in dorsoanteriorized embryos (Kao et al. 1986; Kao and Elinson 1988; Kao and Elinson 1989) (Movie G). This is a major point in establishing the independence of CT from CE, and it is consistent with the idea that CT is expressed early in normal embryos in the same pattern as it is expressed alone and throughout BP closure in ventralized embryos.

In order to determine the pattern of thickening driven by CT, we made standard giant explants from rhodamine dextran injected embryos at stage 9.5 and subsequently fixed them at stage 10 to 10+ or stage 10.5 to 11. These giants were subsequently cut sagittally or parasagittally within the dorsal or ventral portion of the explant and imaged via confocal microscopy, facing the cut surface (Figure 4-supplmentary figure 1C,D). The thickness of the lower and upper IMZ and of the NIMZ were measured in dorsal and ventral sections (Figure 4A). While the expected thinning of the NIMZ was observed, only the lower ventral and upper dorsal IMZ showed substantial thickening over the time period measured. The lack of thickening in the lower dorsal IMZ may be explained by the onset of radial intercalation around stage 10.5 that drives the thinning in the presumptive anterior notochordal region (Wilson and Keller 1991), as part of the transition associated with involution. However, the lack of thickening in the ventral upper IMZ was somewhat surprising. One possibility for the lack of observed thickening is that it occurred prior to or during explantation. We attempt to resolve this issue, below (*Onset of thickening and differential CT in IMZ vs. NIMZ tissue*).

**Figure 4.**
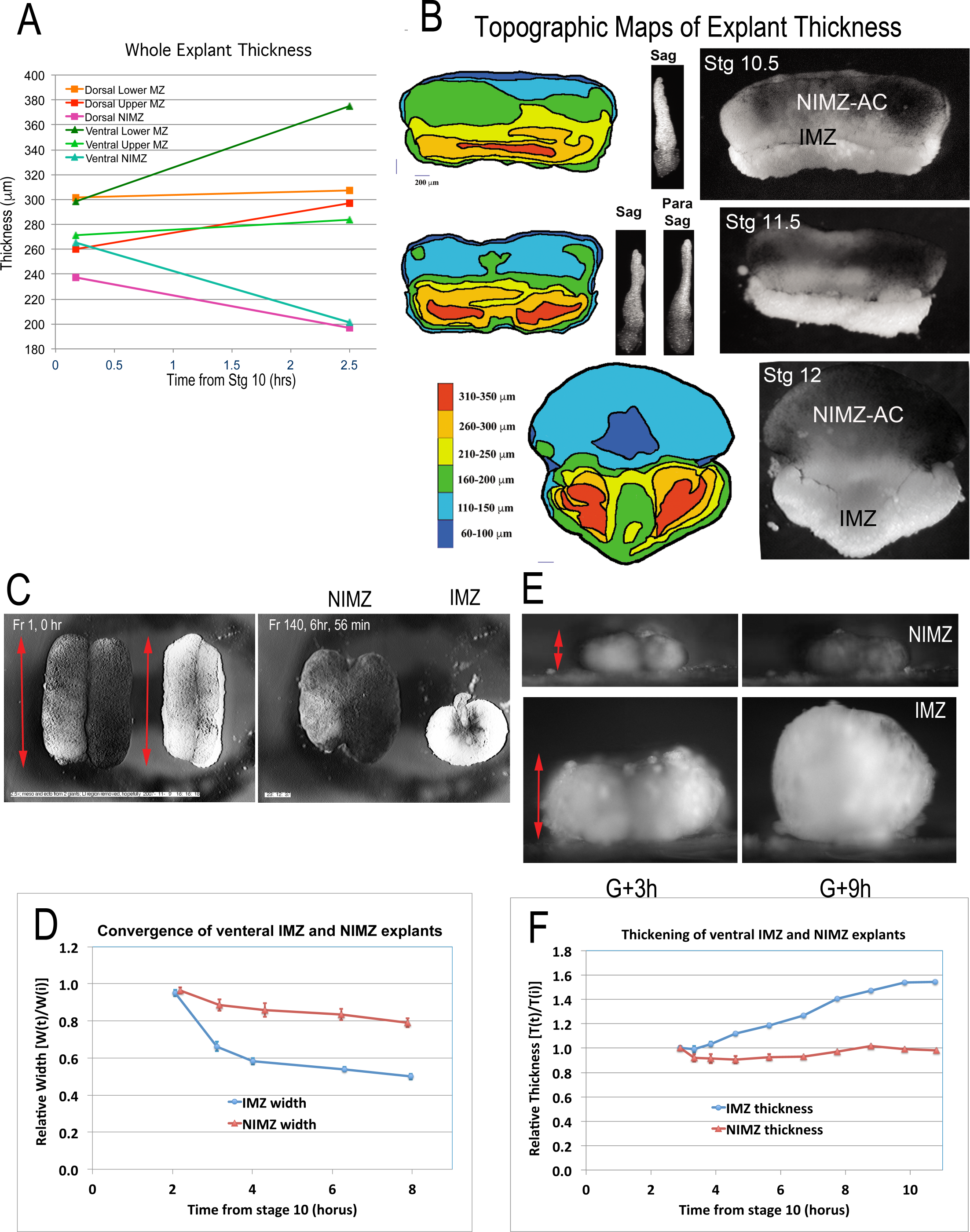
Convergence and thickening in IMZ vs. NIMZ regions. Double NIMZ and IMZ sandwich explants were used to examine the autonomous rates of convergence and thickening in each region, by imaging both from above and from the side, viewing the lateral edge (A, C). Explant width and thickness were measured over time and plotted with respect to the initial value (B, D). The regional thickening of whole explants and epithelial thickness was also determined from sagittal and parasagittal confocal images of Rhodamine dextran injected embryos (E,F; see also Figure 4-supplementary figure 1 C,D). A similar method was used to make rough topographical maps of explants at later stages (G), except that en face confocal images were used. Topographic Maps of regional thickness of explants at 3 stages (left column) corresponding to the time lapse frames (right column) and lateral views (center column).

Later explants were imaged enface to more easily reconstruct topographic maps showing regional variation in thickness (left, Figure 4B). The IMZ continues to thicken in the ventrolateral regions from stage 10.5 to 12, whereas the center section of the IMZ at stage 12 has thinned relative to earlier, which reflects the continuation of thinning by radial intercalation of the notochord and of somitic tissues prior to MIB (Wilson and Keller 1991). Note, however, that the most posterior (animal) portion of the dorsal IMZ remains relatively thick (Figure 4B, yellow arc in Stage 12 topo map). This represents the portion of the dorsal IMZ that has not yet undergone the transitions associated with involution. Whereas much of the thickness of the more anterior lateral and ventral IMZ reflects the attached VE (Figure 4B, right - fluffy tissue at vegetal edge), in the more animal portion of these regions, the thickness represents the continuation of the band of pre-transition material running across the animal edge of the IMZ (Figure 4B, right, orange contiguous with yellow arc). Note also the extreme thinning of the central region of NIMZ at this point, which reflects the center part of the posterior neural ectoderm, which is undergoing CE at this point (see Keller and Danilchik 1988).

### Onset of thickening and differential CT in IMZ vs. NIMZ tissue

The initial identification of the CT movement is based on the observation of persistent convergence and thickening in the IMZ portions of ventral sandwich explants (Keller and Danilchik 1988). However, all tissues, including the NIMZ and AC regions, have some tendency to round up if flattened beyond their equilibrium state (Luu et al. 2011)(other REFs). To establish the extent to which convergence and thickening of the IMZ and NIMZ occurs, to resolve the issue of when such thickening begins, and to determine how convergence and thickening compare in the NIMZ and IMZ, we measured thickness and convergence over time in live explants of NIMZ and IMZ regions. Thickness measures for live explants are less accurate, but this approach allows us to follow the behavior of a single explant over time, removing the issue of embryo-to-embryo variability and variations in explant construction.

To avoid the transition from CT to CE, we made sandwich explants of the ventral 180° of the embryo (V180s) (Figure 1-supplementary figure 1E), which, like explants from ventralized embryos (Figure 1G; Figure 1-supplementary figure 1F) rarely differentiate any dorsal tissues and remain almost entirely ventral in character when isolated before stage 10+ (personal observations and Dale and Slack 1987); thus the CT behaviors can be isolated and more easily studied, over a longer period of time. After allowing the sandwich leaves to heal together, we then cut out NIMZ and IMZ regions, avoiding the interface between the two regions, the limit of involution. To avoid healing artifacts, two NIMZ or IMZ regions were juxtaposed with their newly cut posterior surfaces facing each other and allowed to heal together again as double NIMZ or IMZ sandwich explants (Figure 1-supplementary figure 1H,I; Figure 4C, 0 hours). Prior work has shown that the IMZ converges rapidly during explantation, by roughly 30% of its initial circumference in the intact embryo over the 2.5-3 minutes it takes to cut out the tissue and make a sandwich (Shook et al. 2018); both the IMZ and NIMZ are under strain in the embryo (Beloussov et al. 1990; Chien et al. 2015). Further convergence is slowed or halted when newly constructed sandwich explants are pressed together under a fragment of coverslip glass to allow healing. It is from the point of release from the coverglass that we report convergence (Figure 4D), initially as the two IMZ or NIMZ regions heal together. Double IMZ sandwich explants show more rapid and more extensive convergence than double NIMZ explants (Figure 4C,D), explaining the bowing of giant explants toward the vegetal side. Double IMZs also often loose some or all of their association with the superficial epithelial layer that initially covers them (Movie J).

The convergence of the IMZ in explants described above is accompanied by thickening, seen in movies made from the side (Movie M). In order to estimate how much thickening occurs on initial explantation, we used averages based on histology of whole embryos to give estimates of expected thickness of the IMZ (doubled in Figure 4-supplementary figure 1 B, for ease of comparison) and rough measurements of explants just after explanation. Based on this, IMZs thicken by about 50% during explantation, but then return to roughly their thickness in the intact embryo when the sandwich is pressed together under coverglass. Then, after releasing the sandwich from the glass and cutting out the IMZ regions, explants re-thicken rapidly, by about 50%, while the two regions heal together. NIMZs show much less change in thickness over the course of explantation, sandwich construction and double NIMZ healing. Our reported measures of thickening begin after the two IMZ or NIMZ regions heal together (Figure 4E, G+3h), and are expected to underestimate the amount of thickening of the IMZ, but not the NIMZ.

Double IMZ convergence has an initial rapid phase that correlates with the rapid thickening of the IMZ in the first hour after release from the coverglass noted above (data not shown), then shows a slower phase of convergence. This slower convergence correlates with the continuing, slower thickening of the IMZ. In contrast, the NIMZ never shows rapid mediolateral convergence, and does not thicken (Figure 4E,F), but it does moderately and isotropically elongate in the perpendicular direction within the horizontal plane, i.e. along the animal-vegetal axis (e.g. Figure 4C). For both IMZ and NIMZ regions, there did not appear to be any difference in convergence or thickening in the animal vs. vegetal edges of the tissue. These results demonstrate the distinct properties of IMZ and non-IMZ tissues.

(Experiments to look at convergence and thickening at earlier stages are planned.)

### Onset of other behaviors associated with CT and the transition to CE in explants of the IMZ

The spatiotemporal expression dynamics of CT and its transition to CE is most clearly displayed in explants. The onset of CT is hard to pinpoint, because the IMZ shows convergence from as early as we have been able to measure, after constructing explants and allowing them to heal, and shows no striking increase in the rate of convergence from the onset of measurement (e.g. Figure 3C-E; Movies D-F). There are some clues that suggest it begins with the earliest indications of gastrulation, in particular the onset of apical constriction associated with bottle cell formation at stage 10−. In sandwich explants made prior to stage 10−, wherein the dorsal midlines are identified by “tipping and marking” the embryo during the 1 cell stage, we see that the IMZ and NIMZ regions converge at the same rate when the coverglass is initially removed, up until about stage 10-(G-1h) (working to quantitate data), after which the IMZ consistently converges more rapidly than the NIMZ (Figure 5A, stage 9.5 to 10+; Movies D, Q1). In explants that include most or all of the bottle cell field, we also see the superficial epithelium begin to retract animally at stage 10− as the bottle cells begin to form, coupled with the exposure of the mass of leading edge mesoderm (Movie D). In explants that include less or none of the bottle cell field, superficial epithelium retraction begins later and the deep mesoderm is exposed later (Movie Q). As the deep cells begin to bulge out, they form a mass that retains its integrity but has lost surface contact with the epithelium (Figure 5A, stage 11; Movie Q2). At stage 11, the epithelium begins to re-spread onto this naked mass of deep cells as the bottle cells reverse their apically contracted state, expanding to cover a larger area of the deep cells, and as the epithelial and deep components of the explant re-anneal, they undergo CE (Figure 5A, stage 11 to 12, Movie Q3). In the absence of a cover glass compressing the deep IMZ, which can support CE from stage 10.5 (Shih and Keller 1992a), the re-association of the epithelium with the deep IMZ is required for it to undergo CE (Ninomiya and Winklbauer 2008). In the embryo, the respreading of dorsal bottle cells is associated with the extension of the notochord underneath this epithelium. As CE proceeds, cells are fed out of the thickened collar formed by CT that remains in the animal portion of the IMZ (see previous section, Figure 4B, right, stage 12), and progressively add to the posterior end of the tissues undergoing CE (Shih and Keller 1992b) (Movie D, Q4). This progression of retraction and extrusion behaviors also occur in the IMZ of ventralized giant sandwiches and V180s (Movies E,H), although the formation of bottle cells is delayed by about 2 hours, and respreading of bottle cells may be incomplete and fail to recover much of the mass of extruded tissue.

**Figure 5.**
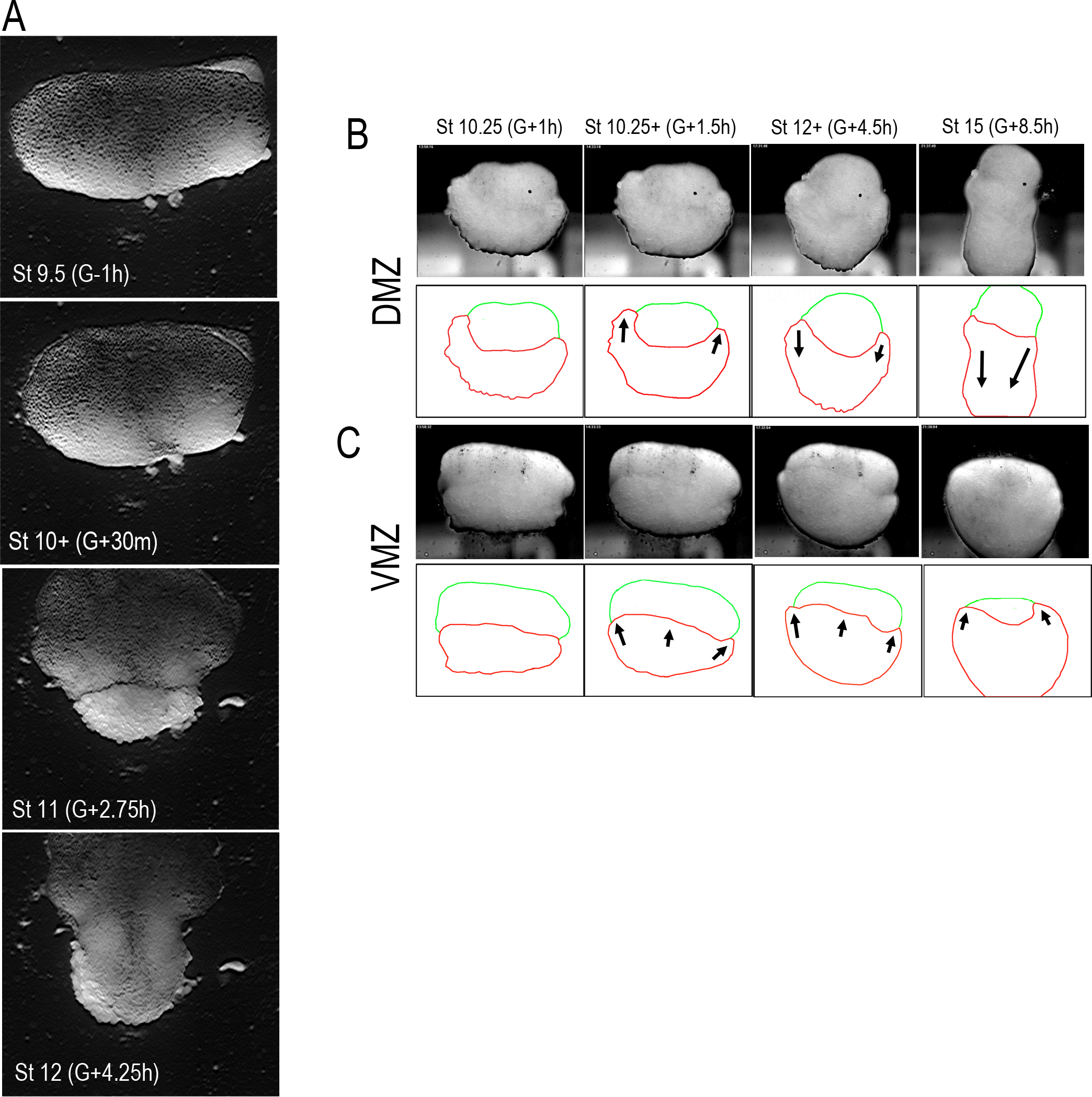
Onset and end of CT. Frames from a time-lapse movie (Movie Q) of a D180 explant show phases in the expression of CT, followed by CE, that reveal their relationship and a potential mechanism for CE (A). Initially (st 9.5 to 10+), the explant converges medially, with greater convergence at the vegetal (IMZ) than animal (NIMZ) and bottle cells form vegetally (the more vegetal bottle cells were not included in this explant). Subsequently, as bottle cells form and the epithelium retracts animally, deep cells are exposed at the vegetal end (stage 10+ to stage 11); in explants with more vegetal IMZ included, this begins as early as stage 10− (see Movie D). CE then begins to extend behind this mass of cells at stage 10.5 and beginning at stage 11, the epithelium begins to respreads dorsally as the epithelium re-anneals with the underlying deep cells (stage 11 to 12.5). This suggests that CT involves a transient loss of adhesion between the deep cells of the IMZ and the overlying epithelium, beginning as early as stage 10−, and that onset of CE involves, and perhaps requires, restoration of a strong deep-epithelial affinity in the IMZ beginning at stage 11; this begins dorsally and progressively sweeps laterally (see Supplemental Movies D, Q). Explants of deep dorsal and ventral upper IMZ plus NIMZ tissue (B, DMZ and C, VMZ) under a coverglass; the upper IMZ (red outline) begins to engulf the NIMZ (green outline) in both DMZ and VMZ explants, but at stage 10.5, IMZ in the DMZ explant reverses course and begins to converge and extend.

An additional behavior associated with the transition from CE to CT is seen in explants of naked dorsal IMZ+NIMZ explants (deep tissue lacking epithelium) under coverglass (Figure 1-supplementary figure 1J). In naked ventral IMZ+NIMZ explants, which never develop dorsal tissues, cells from the IMZ engulf the NIMZ tissue, migrating around the sides of it, and to a lesser extent, over its face, throughout gastrulation (Figure 5C). In contrast, naked dorsal IMZ+NIMZ explants show this behavior until stage 10.5 (when CE begins), then reverse the engulfment and the IMZ begins to converge and extend (Figure 5B). This indicates a change in cell behavior in dorsal tissue, but not ventral; since this coincides with the onset of MIB, it is likely to be associated with a transition to that cell behavior. At this resolution, it is not clear what the deep dorsal IMZ cells are doing; this could be self-isolation as a result of a change in the relative self-affinity (REF) of the dorsal IMZ tissue, or the onset of MIB, or both. CE of dorsal deep tissue compressed under a coverglass from stage 10.5 has previously been observed (Shih and Keller 1992a), but not from stage 10 (Wilson 1990), and activin-induced deep animal cap tissue also shows some ability to converge and extend, even without a coverslip (Green et al. 2004).

These results demonstrate that CT related cell and tissue behaviors begin at about stage 10− and that a second behavioral transition begins at stage 10.5, associated with involution and the onset of MIB. This suggests that as cells involute in the intact embryo, they transition from CT to CE.

### Spreading assays test directly for changes in affinity of the deep mesenchymal and superficial epithelial cells of the IMZ

The expulsion of the deep cells from beneath the epithelial layer in sandwich explants (Figure 5A) suggests a loss of affinity between deep and superficial IMZ tissues during CT. To assay the spreading ability of the epithelial layer on the deep tissue, as a measure of tissue affinity, we made explants from pregastrula stage embryos consisting of deep, dorsal NIMZ+IMZ, which were cultured on fibronectin (FN) substrates. A strip of superficial epithelial tissue was then grafted medially, across the entire animal-vegetal extent of the explant and allowed to adhere under a coverglass (Figure 6A; Figure 1-supplementary figure 1K). The coverglass was then removed, and the area of the strip over the lower (anterior) IMZ, upper (posterior) IMZ and NIMZ regions was recorded over time. Beginning on average around stage 10−, the recombined epithelial tissue lying over the anterior IMZ region began to retract and wad up (Figure 6A, G+2h; Figure 6B; Movie R), in line with our finding, above, that native superficial epithelium begins to retract at stage 10−, and further suggesting that CT-related cell properties begin to change at this stage. Epithelium over other regions remained stable or spread further over time (Figure 6A, G+4.5; 6B). Culture on FN stabilizes the explant by reducing cell motility and preventing it from rolling, thereby making continuous compression under a coverglass fragment unnecessary and allowing greater IFT to develop between the epithelial and deep tissue, in addition to making it much easier to track over time. While FN promotes a small amount of spreading by the deep tissue, especially at later stages, the basic result of epithelial contraction does not change when deep tissue is cultured on a Poly-D-lysine or BSA coated substrate (data not shown). In similar experiments with the deep tissue cultured on a BSA coated substrate and pressed under coverglass to prevent rolling, smaller patches of epithelium contracted and retreated from the surface of the deep IMZ to the NIMZ substrate, where they respread (Figure 5C, Movie S). Epithelial contraction over at least the lower portions of the deep IMZ and spreading over the deep NIMZ or AC tissue occurs regardless of the region of the embryo from which the epithelium was taken (e.g. AC vs. IMZ superficial epithelium), which suggests that it is the deep IMZ tissue that lowers its affinity for the overlying epithelium at the beginning of gastrulation (stage 10, G+0h) rather than the overlying epithelium of the IMZ losing affinity for the deep layer. Unlike the endogenous superficial epithelium that initially lies over the lower, dorsal deep IMZ, recombined IMZ was never observed to respread over this region. These results are a graphic illustration of the loss of affinity between the deep, mesodermal IMZ cells and the overlying epithelium of the IMZ.

**Figure 6.**
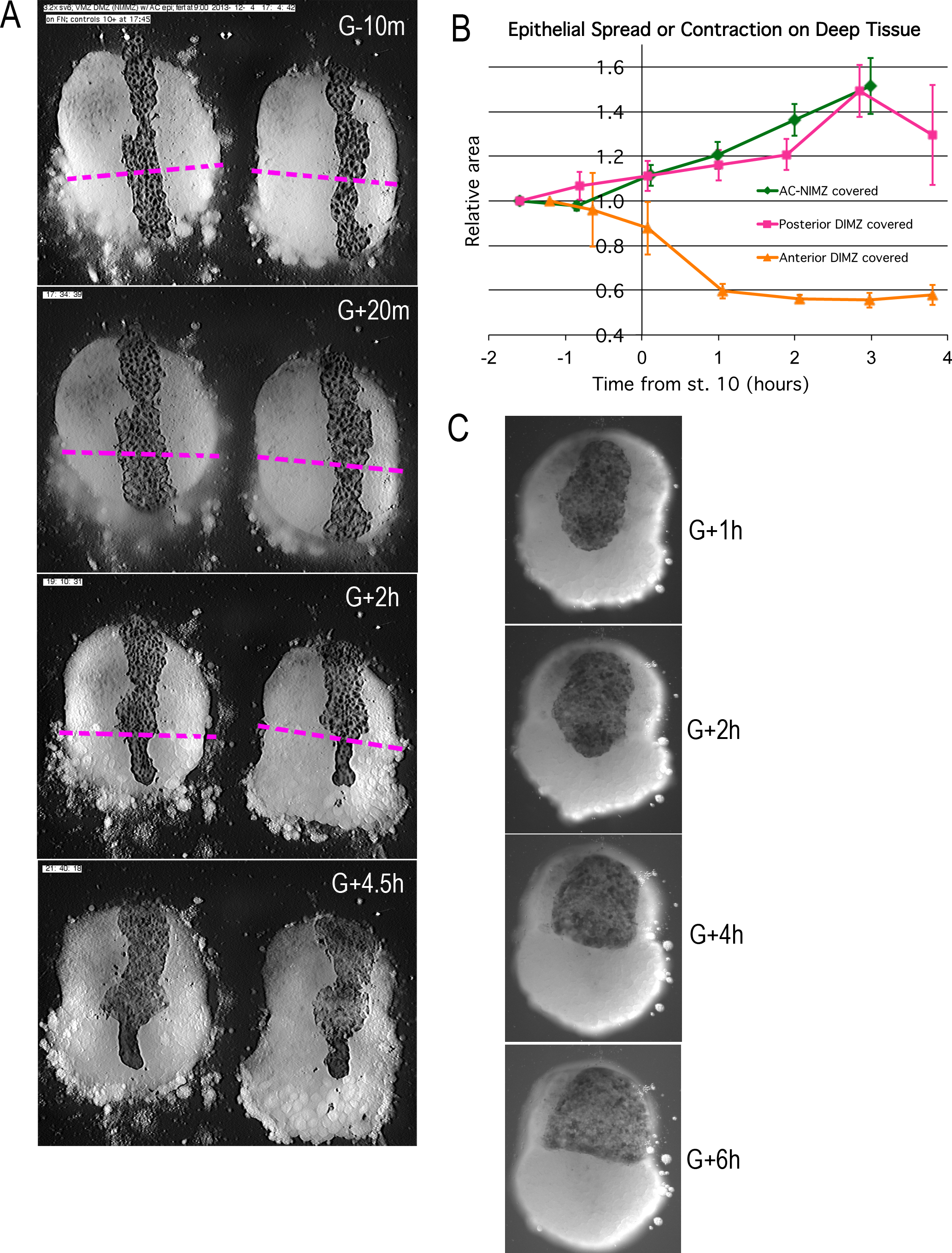
Explants of deep Dorsal IMZ and NIMZ tissue (Anterior DIMZ below dashed line,Posterior DIMZ above, NIMZ further above; boundaries between the three regions identified by cell size and gray cast) made prior to the onset of gastrulation and stabilized by adhering them to fibronectin-coated coverglass; strips of AC epithelium large enough to span the entire animal-vegetal extent of the deep tissue were subsequently grafted on to its surface and kept in place with a coverglass fragment (A,10 minutes prior to the onset of gastrulation (G-10m)). The epithelium adhered and spread over all regions (A, G+20m), at which point the coverglass was removed. At this point, epithelium over NIMZ and posterior IMZ regions contracted slightly, the re-spread, while that over the anterior IMZ contracted more strongly and continuously, often to the point of becoming detached from the deep tissue, and in many cases retracted from the vegetal edge of the deep tissue (A, G+2h). Tissue over the anterior IMZ remained contracted, while that over the posterior IMZ and NIMZ tended to spread (A, G+4.5h).The relative area of spreading or contraction of the epithelial tissue over different deep tissue regions over time was quantitated, beginning after coverglass removal (B). In a variation on this assay, patches of epithelium, lapping across the anterior/posterior IMZ boundary, the epithelium first contracts (C, G+2h), then retracts across the boundary (C, G+4h), where it then respreads (C, G+6h). In this case, the explant remained under pressure from the coverglass pressing the deep and epithelial tissues together throughout the experiment.

### The Interfacial Tension (IFT) Between Deep and Superficial Tissue and the Tissue Surface Tension (TST) of the Deep IMZ increase during CT

Our results showing that the IMZ has a strong tendency to round up (Figure 4), and that at least part of the IMZ reduces its affinity for any superficial epithelial layer, whereas the NIMZ does not, suggested the hypothesis that CT is the result of a change in the TST of the deep IMZ and that this change was driven by a decrease in the affinity between the deep and superficial epithelial tissue, resulting in an increased IFT between the two, driving the observed convergence and thickening in explants as they tend to minimize their surface area. Notably, CE, which tends to increase the surface area-to-volume ratio of the tissue, requires a decrease in the TST experienced by the deep tissue (Ninomiya and Winklbauer 2008), which is in line with the observation that the superficial epithelium respreads over the deep IMZ as it begins to transition from CT to CE. Further, the engulfment behavior seen in naked ventral or dorsal IMZ+NIMZ explants suggests that deep IMZ tissue has a lower TST than that of deep NIMZ (Foty and Steinberg 2005), potentially explaining its lower affinity for the epithelium.

TST is like surface tension of a fluid in that there is a tendency for cells to minimize their free energy by maximizing their adhesive interactions with each other, leading to a tendency to minimize the surface area of the tissue and compact it into a sphere, and in that cells with different affinities (tendencies to form adhesive interactions) for each other are more likely to associate with similar cells, and so self-segregate (Steinberg and Poole 1981; Forgacs et al. 1998). For example, when *Xenopus laevis* tissue fragments are placed onto a flat surface and viewed from the sagittal plane, they round up and form drop-shape aggregates that resemble liquid droplets *in vitro*. In this sense, tissues mimic the behavior of fluids, with a few important differences (Lecuit and Lenne 2007; Green 2008). In particular, TST requires active cell motility in order for cells to rearrange themselves, and the extent of this motility, as well as the cellular cortical tension also play into their affinity for one another, in addition to the kind and number of adhesion proteins they express. For liquids, surface tension and surface energy are numerically identical. Surface energy reflects the work required to expand the surface of a droplet by a unit of area. In other words, surface energy is equal to half the energy required to split a droplet into two equal parts (i.e. to generate two new surfaces); therefore, tissue surface tension is an effective measure of tissue cohesion.

In a morphogenic context, TST can generate additional tension within a tissue if it is in a non-equilibrium state with respect to forces opposing its TST. In the simple case of a clump of tissue or an explant sitting in a dish, TST will generate tension in the most “out-of-round” dimension, causing it to converge and thicken until opposing forces, e.g. from gravity (which acts to flatten the tissue), is in balance with TST. In the whole embryo, TST would generate tension along the longer, circumferential dimension of the IMZ, until it was balanced by the opposing reaction forces from surrounding tissues, such as the bulk of the vegetal endoderm that lies within the hoop of the converging IMZ. Forces arising from more complex sources, such as adhesion to extracellular matrix within the tissue may also counter act TST, e.g. adhesion to FN in AC tissue, helping to drive the thinning and spreading associated with epiboly (e.g. Rozario et al. 2009). Thus in the WE, CT generates tension and convergence but little thickening, whereas in unconstrained explants, CT generates convergence and thickening, but little tension. In constrained explants, CT generates a combination of convergence, thickening and tension (Shook et al. 2018). TST is expected to be the result of unpolarized cell motility, since it is just a matter of cells exchanging contacts to maximize their high-affinity interactions. Polarized force generation then results from geometric context - the surface and shape of the tissue, and any external constraints.

To test the idea that CT was the result of a change in the surface tension of the deep IMZ tissue, e.g. by an increase in the affinity of the deep cells for each other, rather than the epithelium, we measured the TST of explants of deep VMZ and DMZ, using axisymmetric drop shape analysis (ADSA) (David et al. 2009) of deep cell aggregates (Figure 7A-C) by determining the extent to which they flattened under 1 gravity (1g) (Luu et al. 2011). The drop-shape of an aggregate at equilibrium represents a balance of forces between TST and gravity. Since gravity and the densities of *Xenopus laevis* gastrula tissues are known (David et al. 2009), TST can be calculated by the Young-Laplace equation using the radii of the aggregate curvature, which reflects the pressure difference over an interface between two fluids (See Methods and David et al. 2009). The higher the surface tension, the “rounder” the aggregate or “drop” will be at equilibrium. Both VMZ and DMZ deep tissue showed little change in TST until about stage 10, at which point both began to increase through stage 10.5 (2 hours), at which point the DMZ tissue plateaued (Figure 7C), consistent with a transition to MIB. This data is consistent the idea that CT is driven by an increase in deep cell-deep cell affinity, but although our culture media (DFA - see Methods) is designed to mimic the osmolarity, ionic content and pH of interstitial fluid, the deep tissue in the intact embryo is normally in contact with other cells, so our observed results could be an artifactual response of the cells to having a free surface.

**Figure 7.**
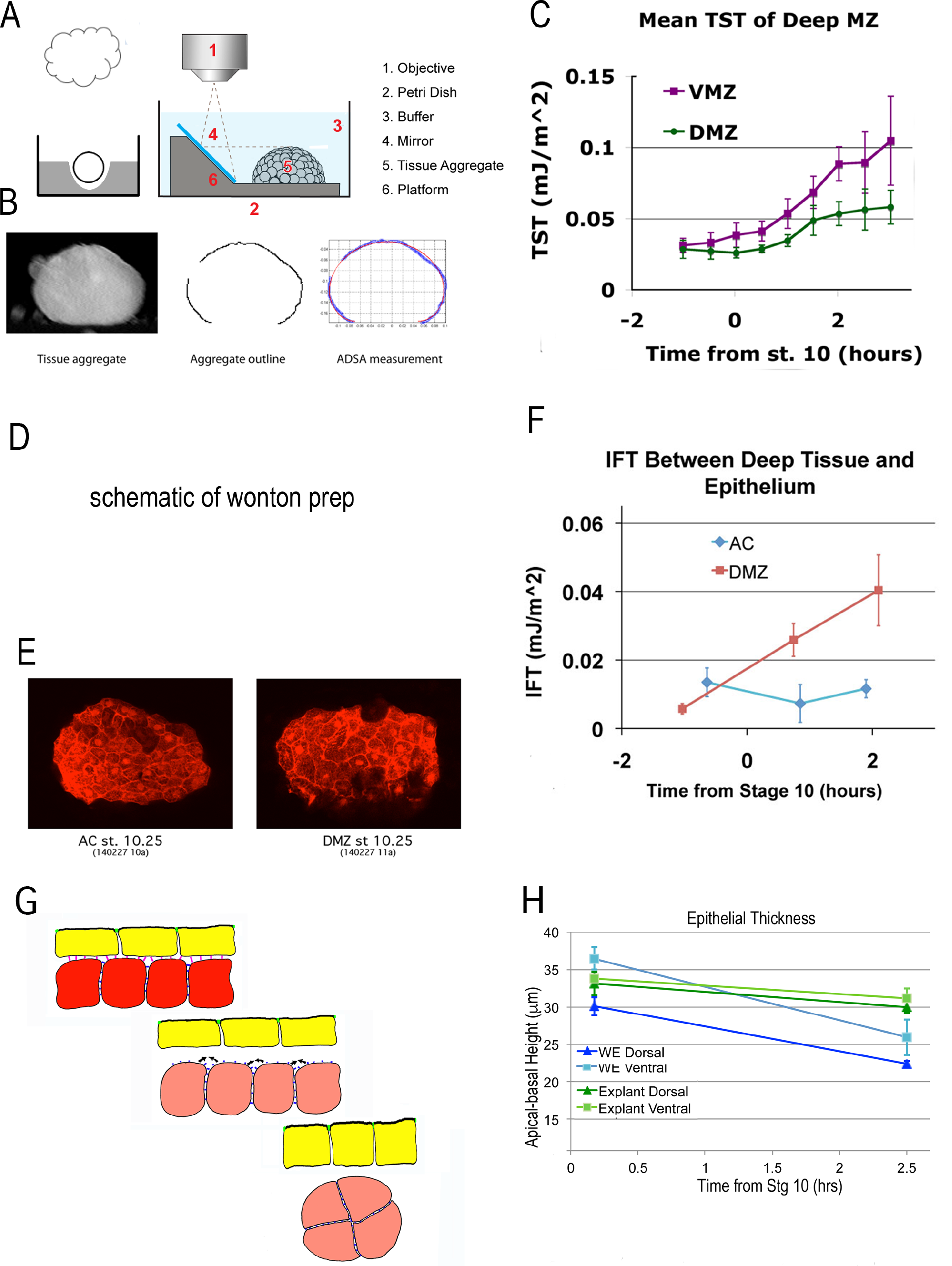
Tissue surface tension and Interfacial tension rise in association with the onset of CT. Deep cell aggregates of IMZ or AC tissue where prepared by removing the superficial epithelium, explanting the deep tissue into a well in an agarose dish, and allowing the explant to round up. (A, left panel). To measure the TST of naked deep tissue in culture media tissues over time, the profile of the aggregate under 1g was imaged using a 45° mirror (A, right panel). Axisymmetric drop shape analysis (ADSA) was used on an outline of the aggregate to determine mean tissue surface tension (David et al. 2009; Luu et al. 2011) (B); mean TST of (n) aggregates was plotted over time for dorsal and ventral IMZ tissue (C, DMZ, VMZ). The mean Interfacial Tension (IFT) between deep tissue and the epithelium was assayed by wrapping aggregates of rhodamine dextran labeled deep tissue in excess unlabeled superficial epithelium (D, the “wonton” explant preparation), letting the explant come to equilibrium and develop for a time, fixing it, cutting the wonton in half and capturing confocal images at 10 μm intervals through the halves. 2-6 confocal slices were projected using max intensity to generate an aggregate outline of the labeled deep tissue (E), which was subjected to ADSA as in (B) to determine the IFT between the deep and superficial tissues at time points from 1 hour before to 2 hours after the onset of gastrulation (St. 10) (F). Model for the development of IFT between superficial and deep layers of the IMZ (G); prior to stage 10−, the superficial epithelium and deep tissue have high affinity for each other, so low IFT; beginning at stage 10−, affinity decreases, increasing IFT and leadingthe deep tissue to round up. One possibility is that the epithelium columnarizes, to match the new, lower surface area of the deep tissue. Epithelial thickness (H) in both whole embryos (WE) and explants declines during the first two hours of gastrulation.

A developmentally programmed drop in the affinity between deep and superficial IMZ tissues would be expected to result in an increase in the IFT between them, as cells within each tissue became more likely to interact with other cells in the same tissue, rather than with those in the other tissue. The consequence would be that the effective TST experienced by the deep tissue would increase, as would its tendency to round up. To measure the IFT, we made deep tissue aggregates from fluorescently labeled embryos, then wrapped them in large pieces of unlabeled superficial epithelium, much like a simple wonton (as in Figure 7D). These were allowed to come to equilibrium for a minimum of two hours under 1g, then fixed at time points. Wonton explants were then cut in half and confocal images collected of the two halves, showing the “drop shape” of the deep tissue filling within its epithelial wrapper (Figure 7E). Control wonton explants, with ectodermal (animal cap, AC) deep tissue filling, did not increase their IFT with respect to the epithelium during gastrulation, but explants with dorsal IMZ (DMZ) deep tissues filling increased their IFT from levels similar to those seen in AC controls prior to the onset of gastrulation, to levels roughly 4 times higher than the AC controls over the first two hours of gastrulation (Figure 7F).

Together these results indicate an increase in deep tissue surface tension (TST) during CT, consistent with an increased cell-cell affinity within the deep mesoderm, and an increase in interfacial surface tension (IFT) between the deep and superficial epithelial tissues in the IMZ consistent with a loss of affinity between the two, but no increase in IFT between the deep and superficial tissues of the ectoderm (animal cap). Both the increase of deep IMZ TST and IMZ IFT would be expected to cause an annular (ring-shaped) IMZ to decrease its surface are, i.e., converge and thicken.

One prediction of such an increase in IFT between the epithelial and deep layers is that there should be increased cortical tension at the cell surfaces facing the interface between the two (REF). However, neither staining for actin nor phospho-myosin light chain (data not shown and Lee and Harland 2007) show an increase in actin localization or myosin activity that would be suggestive of increased cortical tension at this surface.

Since the deep IMZ tissue decreases its surface area by rounding up in explants, one prediction is that the epithelium might also tend to columnarize, thereby decreasing its surface area interacting with the deep tissue (Figure 7G). To test this, we measured the heights of IMZ epithelial cells in whole embryos as a control, and in explants between the onset of gastrulation and the onset of MIB. However, explant epithelia was initially about the same as that in intact embryos, indicating that it had not thickened on explantation, and epithelia further thinned by 5-10% (Figure 7H), suggesting that epithelial “slack” is taken up by some other mechanism. One possibility is the apically contracting, retracting epithelium, which we did not measure.

In an attempt to validate the idea that the observed ventral TST (σ_em_ = 0.08 mJ/m^2^, by G+2h) could be responsible for generating the measured forces generated by CT (about 0.3 μN by G+2h) (Shook et al. 2018), we calculated the expected force of shortening along the long axis as

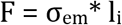

where l_i_ = the length of the contour of the interface between epithelial and deep tissue in a cross section orthologous to their long axis, which averages about 1.5 mm, such that F = 0.12 μN, or a factor of about 40% of the observed force. Using the dorsal IMZ TST or IFT yields a lower number. If these calculations are correct, this suggests that some other mechanism may be involved in generating about 60% of the observed convergence force during the first two hours of gastrulation. One possibility is the bottle cells, which account for a noticeable delay in force increase when they re-expand during tractor pull assays (Shook et al. 2018). Another possibility is that the deep cells are engaged in a different, or additional type of motility, associated with the convergence required for involution (e.g. Evren et al. 2014).

(Additional measurements and analysis underway to further validate or refute these results; statistical analysis of IFT and interface contour measures; plate compression measures of TST in IMZ vs. NIMZ; compressive (thickening) force generation by double IMZ explants undergoing CT)

### CT relies on different mechanisms of cell motility than CE to generate force

To test the idea that CT is dependent on a cell motility distinct from that driving CE (MIB), with differential molecular control and effectors, we disrupted two molecules shown to be required for MIB, one involved in force generation and one in patterning MIB, with the expectation that normal embryos, which depend on a combination of CT and MIB to generate force for blastopore closure, would be more affected than ventralized embryos, which depend only on CT. An underlying assumption here is that MIB is not a process layered on top of CT, but one that represents a transition to a fundamentally distinct mode of motility; thus if elements of MIB are broken, but cells still make the transition from CT to CE, that population of cells will neither continue to generate force by CT, nor effectively generate force by MIB.

Myosin heavy chain IIB (MHC IIB) is up-regulated strongly in dorsal tissues expressing CE (Bhatia-Dey et al. 1998), and a morpholino knock down of MHC IIB strongly inhibits blastopore closure in normal embryos that express CE (Skoglund et al. 2008) (Figure 8A, compare first to second column), whereas the same dose of MHC IIB MO only mildly retards blastopore closure in ventralized embryos (Figure 8A, compare third to fourth column). Only MHC IIB MO injected normal embryos showed a significant decline in frequency of blastopore closure (Figure 8B; p < 0.001 compared to uninjected controls or MHC II B MO injected ventralized embryos). This suggests that CE is strongly dependent on MHC II B, whereas CT is not.

**Figure 8.**
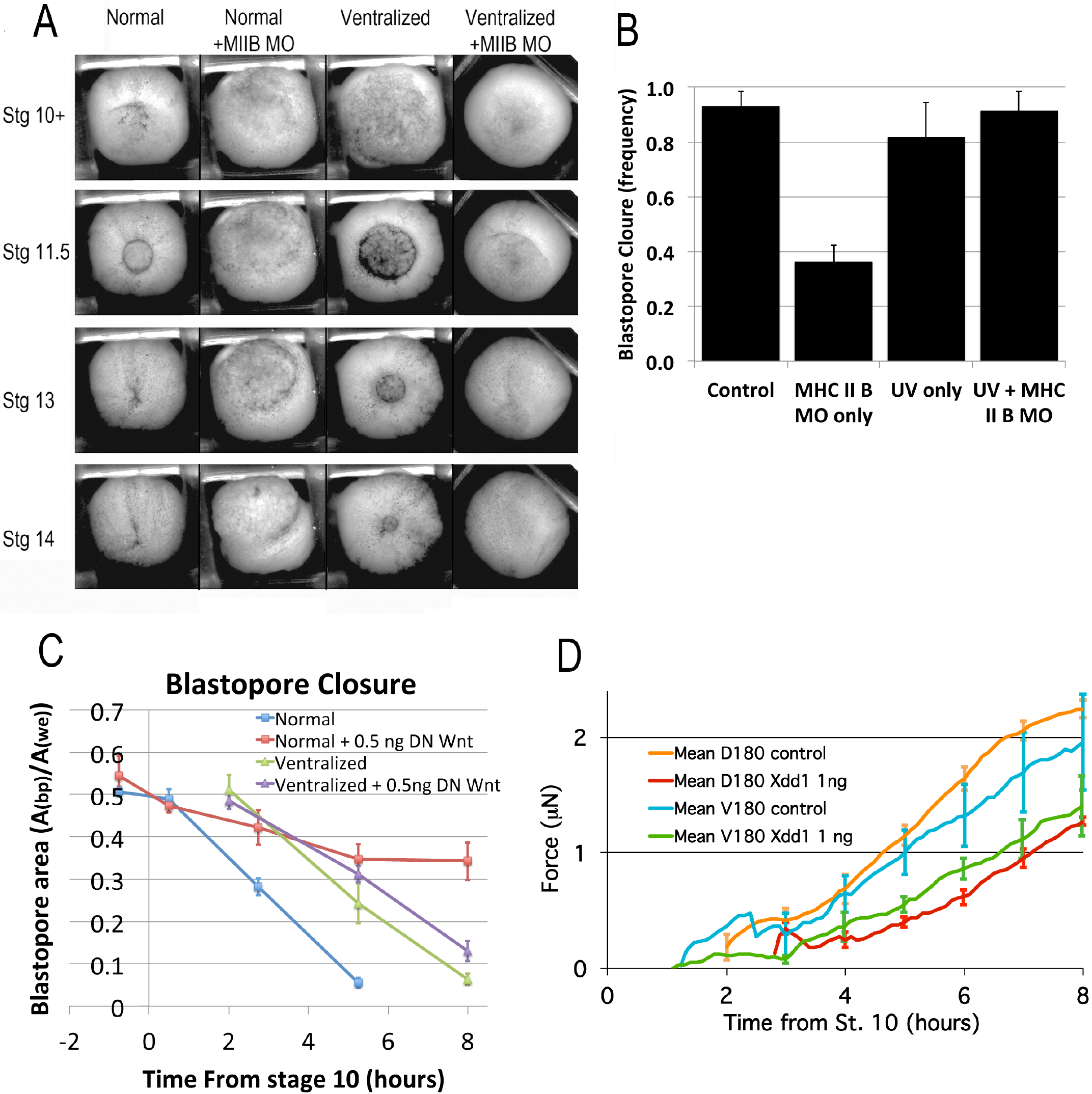
CT and CE depend on different molecular pathways. Representative stills (A) of time lapse movies (Movie B) comparing blastopore closure in normal and ventralized embryos, uninjected or injected with an MHCIIB morpholino. Frequency of blastopore closure by stage 20 was collected for 31 to 90 embryos in each class from 4 different clutches of embryos (B). Comparison of extent of blastopore closure over time in normal and ventralized embryos that were either uninjected or injected with 0.5 ng dnWnt11 RNA (C), based on tracings of the projected area of the blastopore divided by the projected area of the whole embryo, measured from time lapse movies (for example, see Movie C). Force measurements (Shook et al. 2018) of D180 or V180 explants made from embryos that were either uninjected or injected with 01 ng Xdd1 RNA (D).

Disruption of the non-canonical Wnt/PCP signaling pathway strongly inhibits CE in vertebrates, and in particular, disrupts MIB in Xenopus (Wallingford et al. 2000), and interferes with blastopore closure in normal Xenopus embryos (Sokol 1996; Tada and Smith 2000). One prediction of the hypothesis that CT is the result of tissue surface tension (TST) is that since the cell motility underlying TST is not polarized, disrupting the planar cell polarity (PCP) pathway should have less of an effect on tissue movement and force generation driven by CT than by CE; this is supported by previous work showing that embryos injected with Xdd1 RNA dorsally, but not ventrally suffer blastopore closure delays (Ewald et al. 2004).

To test this idea, we compared blastopore closure in normal and ventralized embryos, that were either uninjected or injected with 0.5 ng dnWnt11 RNA (Tada and Smith 2000). Whereas dnWnt11 RNA had the expected result of slowing and often completely stalling blastopore closure in normal embryos, it caused only a minor delay in BP closure in ventralized embryos (Figure 8C; Figure 8-supplementary figure 1; Movie C), which do not express CE. To show that force generation by CT is also primarily dependent on mechanisms other than those blocked by PCP disruption, we compared forces generated by Dorsal 180° (D180) and V180 sandwich explants from embryos that were either uninjected or injected with 1 ng Xdd1 RNA (Sokol 1996). Xdd1 RNA injection strongly decreased the amount force generated by D180 sandwich explants (by about 63% by the end of gastrulation at G+6h - compare red and orange force traces, Figure 8D), which express CT early but then show the typical progressive transition to CE, whereas Xdd1 overexpression has less effect on force generated by a V180 explants (by about 35% at G+6h; compare green and blue force traces, Figure 8D), which express only CT. This indicates that force production by CT does not depend on PCP signaling as strongly as CE does. That Xdd1 injected D180s generate only about 40% as much force as uninjected V180s supports the conclusion that the D180s do not continue to express CT when MIB motility is blocked. (Additional tractor pull tests underway to improve n’s)

## METHODS

### Embryo manipulations & microinjection

*X. laevis* embryos were obtained and cultured by standard methods (Kay and Peng 1991), staged according to Nieuwkoop and Faber (Nieuwkoop and Faber 1967), and cultured in 1/3X MBS (Modified Barth’s Saline).

To ventralized embryos, de-jellied embryos were placed in dishes made of 15 mm transverse sections of 60 mm diameter PVC pipe with Saran wrap stretched across the bottom, irradiated 6 or 7 minutes from below at about 35 minutes post fertilization on a UV trans-illuminator (analytical setting, Fotodyne Inc. Model 3-3500) and left undisturbed for at least an hour to avoid accidentally rotating and thus dorsalizing them (Black and Gerhart 1986). Embryos forming bottle cells asymmetrically or earlier than the majority of ventralized embryos were discarded as being insufficiently ventralized. To dorsoanteriorize embryos, embryos were exposed briefly to Lithium chloride (0.35 M in 1/3X MBS at the 32 cell stage, for 6 minutes)(Kao et al. 1986). Control, ventralized or dorsoanteriorized embryos were cultured to control stage 35-38 and scored for their DAI (Kao and Elinson 1988) to evaluate the effectiveness of ventralization or dorsalization.

To label with rhodamine dextran amine (RED), embryos were injected at the 1-2 cells stage with 25-50 ng RDA (Molecular Probes, D1817) and fixed at the desired stages with MEMFA (Kay and Peng 1991). To knock down MHCIIB, embryos were injected with 7.5 pmol MO (Skoglund et al. 2008) per embryo. To disrupt PCP signaling, embryos were injected with 1 ng Xdd1 RNA (Sokol 1996) or 0.5 ng dnWnt11 (Tada and Smith 2000). Xdd1 RNA efficacy in the embryos used for the tractor pull was confirmed by an absence of CE in dorsal tissue, either in the sandwich explant in the tractor pull, or tissue removed from the explant, for D180 and V180 explants, respectively.

### Explant preparations

Explants are cultured in Danilchik’s for Amy (DFA) (Sater et al. 1993). For explants made before stage 10 the embryos were tipped and marked to identify the dorsal side (Sive et al. 2000). Standard “giant” sandwich explants were made at stage 10 to 10.25 with bottle cells and some vegetal endoderm included (Shook et al. 2004), or without bottle cells (Poznanski et al. 1997); ventral giant sandwich explants were made as above, but initially cutting along the dorsal, rather than ventral midline (REF?) (see also Figure 1-supplementary figure 1A-C). Dorsal 180°, and Ventral 180° explants are made as standard or ventral giants, with the ventral or dorsal 180° of the marginal zone cut off (Figure 1-supplementary figure 1D,E). Ventralized or dorsoanteriorized giant explants were made from ventralized or dorsoanteriorized embryos (Figure 1-supplementary figure 1F,G) without reference to “dorsal” as these embryos are symmetrical about the blastopore (Scharf and Gerhart 1980; Kao and Elinson 1988).

Double ventral or dorsal, IMZ or NIMZ only sandwich explants (Figure 1-supplementary figure 1H,I) were made by first constructing V180 or D180 explants, allowing them to heal together for a minimum of 30 minutes, then cutting a few cells above or below the limit of involution (LI), and recombining two NIMZ or IMZ pieces with their newly cut posterior edge juxtaposed and pressed together by two coverglass bits on either side of the combined explants; the double explants were allowed to heal for 1 hour before being moved onto a coverglass bit platform in front of a 45° mirror in the culture dish, with the lateral edge of the explant facing the mirror, such that both convergence and thickening could be imaged.

Naked dorsal or ventral IMZ+NIMZ explants (Figure 1-supplementary figure 1J) were made by peeling the superficial epithelium off the dorsal or ventral-most ~90° of the embryo at stage 9 to 10, cutting the peeled sector out of the embryo, trimming away vegetal endoderm, and pressing the remaining tissue under coverglass, allowing observation of the lateral engulfment behavior. In some cases, the explant was covered with a thin sheet of agarose in between the surface being imaged and the glass, to reducing sticking and observe more native cell motility. For epithelial spreading assays, the inner face of the explant was pressed against FN coated glass to which it rapidly adhered, and a strip of superficial epithelium was laid along the animal to vegetal extent of the explant (Figure 1-supplementary figure 1K). The epithelium was held in place with a coverglass to prevent it from curling up on itself and to promote adhesion with the deep tissue; after about 30 minutes, the glass was removed.

To make deep tissue aggregates, superficial epithelium was removed from AC, dorsal or ventral IMZ regions at stage 9-9.5, and the exposed deep tissue cut out and trimmed down to make sure it included no other tissue type. Clumps of deep tissue were then placed in agarose wells to promote aggregation for 30-60 minutes. Aggregates were then placed on coverglass platforms in front of 45° mirrors and imaged from the side to observe rounding over time. To make wontons, such aggregates made from RDA labeled deep tissue were instead wrapped in excess epithelium from unlabeled embryos, such that the edges of the epithelium healed together all around the deep tissue; in some cases, these were also imaged from the side. Wontons were allowed to age to specific
control stages, then fixed in MEMFA, cut through the middle, cleared, positioned with their cut surface down, and imaged via confocal.

### Imaging

Explants via dissecting scope (& w/ mirror)

Explants via inverted

Explants via upright w/ dipping lens

NIH image, Scion Image, ImageJ, Metamorph

### Confocal Microscopy

RDA explants

### Morphometics & image processing

Object Image, ImageJ

### Surface Tension Measurements

Tissue fragments excised from the early gastrula (stage 10) were placed in a well for 30 minutes to facilitate rounding. Tissue aggregates were transferred onto tissue culture substrates pretreated with non-adhesive coating and allowed to equilibrate for 2 hours before aggregate profiles were captured using an inclined mirror calibrated at a 45° angle to the substrate surface for imaging. Tissue surface tension was quantified using the Axisymmetric Drop Shape Analysis (ADSA) algorithm (Rio and Neumann 1997) modified for use with tissues by introducing more robust image parameters to accommodate the irregular curvatures of aggregates (David et al. 2009; Luu et al. 2011; David et al. 2014). The curvature outline was identified using an edge detection algorithm (Canny) in MATLAB. All processed images were inspected prior to measurements to ensure that Canny detections generated accurate depictions of the tissue aggregate. ADSA is a program written for MATLAB that numerically integrates the Laplace equation to generate theoretical drop-shapes for hypothetical surface tensions, which are optimally fitted onto experimental profiles obtained from tissue aggregates. The drop-shape that best fits the aggregate is used to derive the specific tissue surface tension for that particular aggregate.

The Young-Laplace equation of capillarity defines the equilibrium of a liquid surface as the following:

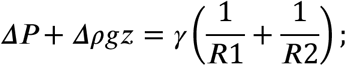

from left to right, ***ΔP*** is the difference in pressure across the interface between the tissue and the immersion medium, ***Δρ*** is the difference in density between the tissue aggregate and the immersion medium, *g* represents gravitational acceleration, *z* is the vertical distance from the aggregate apex, *γ* is the tissue surface tension, and *R*_1_, *R*_2_ are the radii of the aggregate profile.

### Tractor pull force measurements

Forces generated along the mediolateral axis by sandwich explants were assayed using the tractor pull apparatus (Shook et al. 2018). More recent experiments were done using 250 μm beads, as they generate less friction with the sled than 100 μm beads.

## DISCUSSION

These results constitute a major revision of how amphibian gastrulation occurs and establishes CT as a bona fide morphogenic movement (Figure 1).

Closure of blastopores in ventralized embryos was first reported by Scharf and Gerhart (1980), a classic paper that stimulated the current work. These results indicate that there is another force-generating morphogenic machine functioning to close the blastopore, and that it can, in fact, close the blastopore alone, without CE.

### Onset of CT

Bottle cell retraction & loss of epithelial affinity

Increase in VMZ and DMZ TST

Increase IFT w.r.t. epithelium of DMZ deep compared to AC deep

Differential IMZ/NIMZ convergence and thickening (is there a point before they become different?)

### Transitions from CT to CE

Expression of RI prior to and MIB during involution (Wilson and Keller 1991; Shih and Keller 1992a); relation to a requirement for FN (Marsden and DeSimone 2001; Marsden and DeSimone 2003).

Bottle cell re-spreading & surface tension requirement for CE (Ninomiya and Winklbauer 2008): Regaining affinity between these layers and reduction of tissue surface tension has been shown necessary for CE to occur effectively as it involves an increase in tissue surface area relative to volume.

This explains the common observation by embryologists that the surface epithelium of the pre-involution IMZ is particularly easy to remove from the underlying deep layer but more difficult after the IMZ involutes.

### Potential role for TST driving CT

Mechanism of TST & caveats to causes (DAH vs. cortical tension; Green) TST in morphogenesis more broadly; Holtfreter; Steinberg & Takeichi; Brodland; Heisenberg

### TST as Mechanism of CT

Many of the behaviors we document in explants are consistent with CT of the IMZ being driven by TST. IMZ explants released from compression under coverglass rapidly round up for the first 2 hours, indicating that they have been released from an equilibrium state and suddenly find their TST far from equilibrium with external forces. NIMZ explants show much less rounding behavior, indicating they are much closer to equilibrium, and are much closer to equilibrium in a flattened state. Measurements show that both the TST and IFT of IMZ tissue increases from the onset of gastrulation, while that of NIMZ tissue remains constant. The observed changes in deep-epithelial affinity are also consistent with an increase in deep-superficial IFT in the IMZ, but not the NIMZ.

Increase in the IFT between deep and superficial IMZ layers could result from any change in either layer that reduced the affinity, or tendency to form bonds, of one layer for the other. This could result from an increase in the relative affinity of cells within either or both layers for other cells within the same layer, resulting in a decrease in their relative affinity for cells in the other layer. And changes in affinity can result from changes in adhesion, in cortical tension or in motility. Thus, it is not simple to resolve what drives the increase in IFT between the two layers.

Our measurements of TST and IFT suggest that these mechanisms alone may not be enough to explain the convergence forces generated by IMZ tissues. Additional sources of tension generation could come from the epithelium, e.g. by contraction, or other cell behaviors of the deep layer. Potential alternative/ additional mechanisms of CT (Evren).

### Model incorporating CT into gastrulation

This summary resolves many paradoxes in our understanding of gastrulation, reveals for the first time a new and cryptic morphogenic movement, CT, acting in parallel with CE (Figure 9). This explains the fast formation of the mysterious thickened region in the initial behavior of explants, a thickened region which does not exist, at least not to the same extent in vivo, the difference being due to the resistance to CT in the embryo, which is lacking in explants. More important, this work strongly suggests that a change in tissue affinity and the resulting change in tissue surface tension, at least in part, drives CT. It may not be the only mechanism but it is likely a major component. In the past, this type of change or difference in tissue affinity/surface tension was used to explain tissue and cell sorting out, and ordering of layers in the Steinberg DAH (Differential Adhesion Hypothesis) (Steinberg 1970; Steinberg and Takeichi 1994; Steinberg 2007), a hypothesis that was more successful in being consistent with major features of germ layer and tissue type arrangements, particularly in cultured tissues (e.g. Townes and Holtfreter 1955), but less so in explaining details of morphogenesis. What appears to be happening in this case, is a change in tissue affinity in a particular geometry, and annulus of tissue, which results in a circumferential convergence, a “squeezing” ring, by virtue of the initial shape of the tissue, or in the case of the explants, a linear, largely mediolateral convergence force which we have measured (see Shook et al. 2018). This raises the question of how many other examples there are of patterned, oriented forces arising from changes in a general property, tissue affinity and surface tension, in specific geometries and with specific anchorage points.

**Figure 9.**
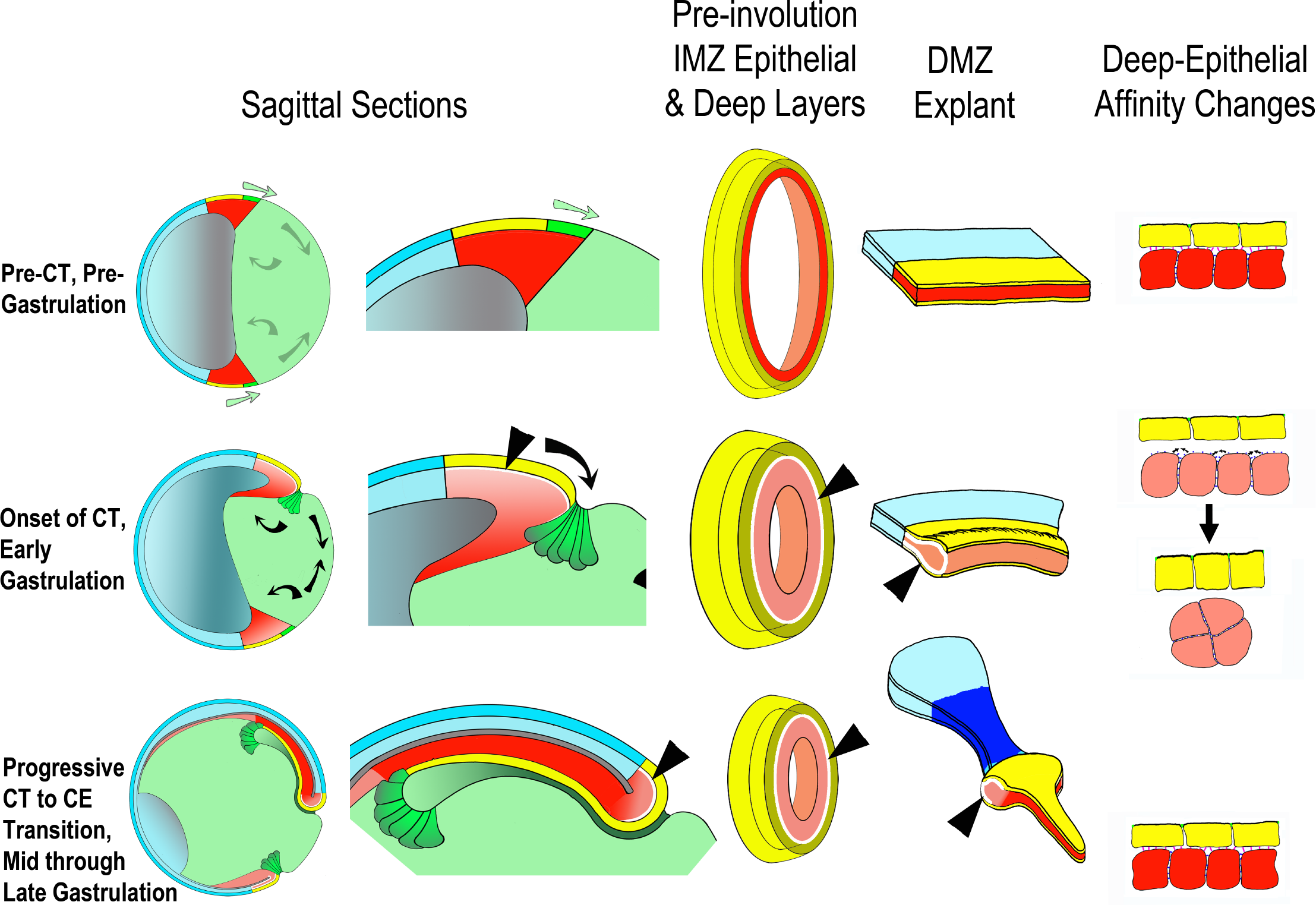
Proposed mechanism of CT, its function in the gastrula and in explants, and its relationship to CE. Prior to the onset of gastrulation (top row), vegetal rotation (gray arrows, Sagittal Sections) has not yet occurred and the IMZ annulus remains stable, as does the IMZ explant, due to a stable deep-epithelial cell affinity (far right). With the onset of gastrulation and CT (second row), vegetal rotation, apical constriction of the lower IMZ epithelium (bottle cell formation), involution and invagination also begin (black arrows, Sagittal Sections). At this time, the deep mesodermal tissue begins to loose its affinity with the overlying epithelium; initially this occurs along the vegetal edge as the bottle cells form, but by stage 10.5 both surface tension of the IMZ as a whole and its interfacial tension with an epithelium increases, resulting in a tendency for the IMZ to minimize its surface area by thickening (Second Row, far right). This loss of affinity at the interface of deep and epithelial tissue (shown by white spaces between epithelial and deep, at black arrowhead pointers) results in thickening of the IMZ of the explants, and would tend to thicken and decrease the circumference of the IMZ annulus, thus generating a preinvolution force that tends to converge the IMZ around the blastopore (center column, second row). Although the force is generated, in the intact embryo little thickening of the IMZ occurs, both because it is resisted by adjacent tissues (vegetal endoderm, ectoderm), and because in the embryo, convergence is coupled with involution, progressively removing the material of the IMZ. During early involution, as the lower edge of the IMZ involutes, it transitions from expressing CT to expressing directed migration (second column, second row), and during later involution, the upper IMZ transitions to expressing CE (second column, third row). In the case of CE, this transition is accompanied by regaining a high deep to epithelial affinity (loss of white space, on involution, second column, second and third rows). In the explant, the convergence force is not resisted by the vegetal endoderm, nor does the lower IMZ migrate away and so convergence generates a thickened collar (CT, second row, bulge in DMZ explant). At stage 10.5 (G+2h), tissue laying dorsally at the vegetal edge of the upper IMZ begins to transition to expressing CE in an anterior to posterior progression (Shih and Keller, 1992) and extends the dorsal axial and paraxial tissues from the thickened collar, pushing the leading edge mesoderm (not currently shown in figure) in front of it and to the sides. In the meantime, Vegetal Rotation (black arrows, first column, second row) begins to change the shape of the vegetal endoderm at the onset of gastrulation such that convergence force generated by CT cooperates with VR to move the IMZ vegetally and push the vegetal endoderm inside. When CE begins at stage 10.5, it has been positioned by the actions of CT and VR such that it further moves the IMZ vegetally and pushes the vegetal endoderm inside, in contrast to converging in its initially more equatorial position, where it would tend to push the vegetal endoderm out, rather than in. The convergence driven by CT and CE also pulls the ectodermal (NIMZ-AC) region vegetally. Finally, the structural liability of the relatively weak attachment of the IMZ epithelium to the underlying deep region, implied by the apparent low affinity or adhesion between them, is offset by the fact that in the intact embryo, the IMZ epithelium is tightly attached at its vegetal edge to the forming bottle cells, which in turn are tightly attached to the adjacent vegetal endodermal tissues. Thus it is pulled along with them as they move inside the blastopore.

### Evolutionary conservation of CT

Conservation of CT in anuran BP closure and addition of CE at different times among anurans.

Broader conservation of CT as a mechanism for blastopore closure in single layered epithelium (Wolpert as an example of mechanism, (Luu et al. 2011) as a potential example of epithelial thickening)

### Discriminating the causes of BP closure (or just how you could tell if it was a defect in CT?)

Failure due to defects in CE vs. CT vs. VR vs. epiboly (Involution, tissue separation, mesendoderm migration)

### Conclusion

We have demonstrated the function and mechanism of CT and shown that it is capable of generating at least some of the mediolateral convergence forces of the IMZ.

## ACKNOWLEDGEMENTS

NICHD R37 HD025594 MERIT AWARD to Ray Keller

NIH RO1 GM099108 to Paul Skoglund

---

**Figure 1-supplementary figure 1.** Description of explant construction. Lateral view of the pre-gastrulation embryo (st. 9.5) showing presumptive germ layers overlaid on AC, NIMZ, IMZ and VE, with region of bottle cell formation and Limit of involution (LI) indicated (Z). <u>Standard “giant”</u> sandwich explants were made at stage 10 to 10.25 with bottle cells and a small amount of vegetal endoderm included (A), or with the bottle cell forming field cut off (B); <u>ventral giant</u> sandwich explants were made as standard giants, but initially cutting along the dorsal, rather than ventral midline (C). <u>Dorsal 180°</u> (D180, D), and <u>Ventral 180°</u> (V180, E) explants are made as standard or ventral giants, with the ventral or dorsal 180° of the marginal zone cut off. <u>Ventralized</u> (F) or <u>dorsoanteriorized</u> (G) giant explants were made as above (A) from ventralized or dorsoanteriorized embryos, but without reference to “dorsal”, as these embryos are symmetrical about the blastopore. <u>Double ventral or dorsal, IMZ</u> (H) <u>or NIMZ</u> (I) only sandwich explants were made by first constructing two V180 or D180 explants, allowing them to heal together for a minimum of 30 minutes, then cutting a few cells above and below the limit of involution (LI), and recombining the two NIMZ and the two IMZ pieces with their newly cut posterior edges juxtaposed and pressed together by two coverglass bits against the anterior surfaces of the individual pieces. The double explants were allowed to heal for 1 hour before being moved onto a coverglass bit platform in front of a 45° mirror in the culture dish, with the lateral edge of the explant facing the mirror, such that both convergence and thickening could be imaged. <u>Naked dorsal or ventral IMZ+NIMZ</u> explants (J) were made by peeling the superficial epithelium off the dorsal or ventral-most 90° of the embryo at stage 9 to 10, cutting the peeled sector out of the embryo, trimming away vegetal endoderm, and pressing the remaining tissue under coverglass, allowing observation of the lateral engulfment behavior. In some cases, the explant was covered with a thin sheet of agarose in between the surface being imaged and the glass, to reducing sticking and observe more native cell motility. For <u>epithelial spreading tests</u>, the inner face of the naked IMZ+NIMZ explant was pressed against FN coated glass to which it rapidly adhered, and a strip of superficial epithelium was laid along the animal to vegetal extent of the outer face of the explant (K). The epithelium was held in place with a coverglass to prevent it from curling up on itself and to promote adhesion with the deep tissue; after about 30 minutes, the glass was removed.

**Figure 3-Supplementary Figure 1.**
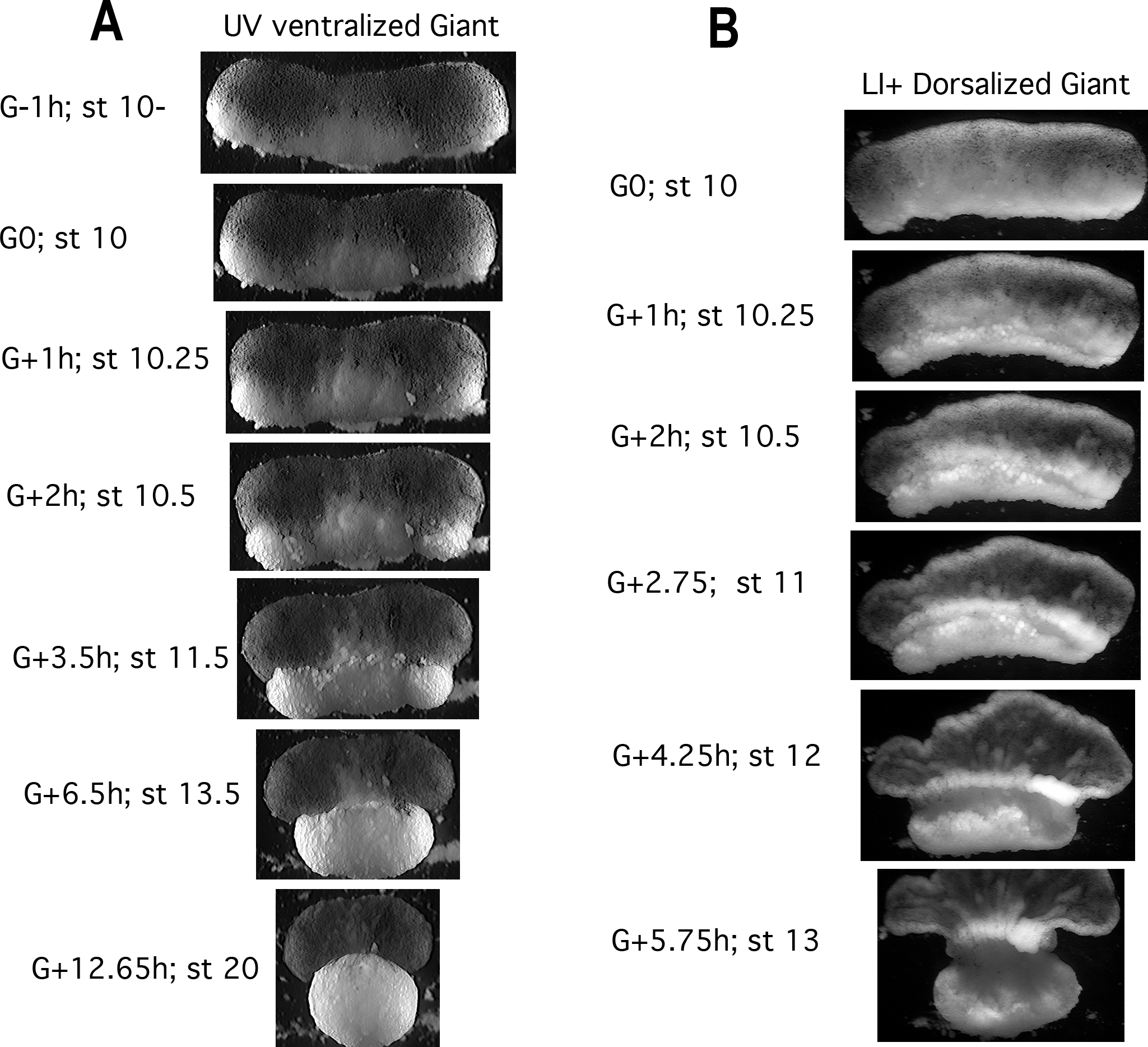
CT in UV Ventralized and Li+ Dorsoanteriorized giant sandwich explants. Giants made from ventralized embryos (A) show isotropic convergence and thickening of the IMZ throughout gastrulation and neurulation in control embryos (stages and hours from the onset of gastrulation (G+Xh) indicated). Giants made from dorsoanteriorized embryos show isotropic convergence and thickening of the IMZ through at least stage 10.5, when they begin isotropic convergence driving anterior-posterior extension; at this point a thickened mass remains in the more animal portion of the IMZ, which eventually feeds into the converging and extending tissue.

**Figure 4-supplementary figure 1.**
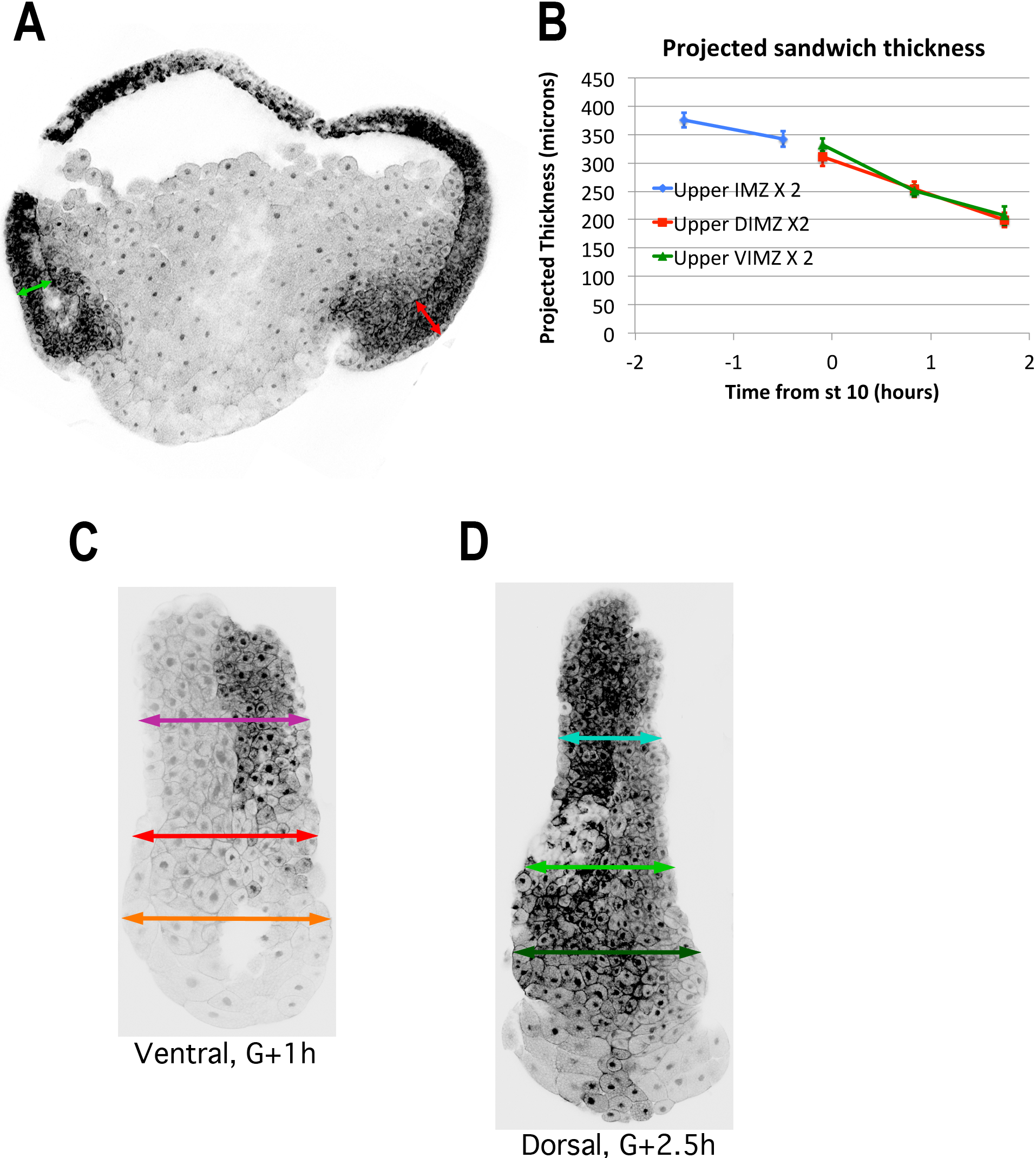
Confocal images of rhodamine dextran injected embryos and explants made from such embryos were used to measure the thickness of the IMZ and NIMZ, and of the superficial epithelium over the IMZ, before and during early gastrulation. In gastrula stage embryos, the thickness of the upper (more animal) IMZ was measured dorsally and ventrally (A, red and green double-headed arrows, respectively). The thickness of the lower IMZ in whole embryos after the onset of gastrulation is more ambiguous, so is not reported. In pre-gastrula embryos, upper IMZ thickness was measured on opposite sides and averaged (not shown). The expected thickness of the resulting sandwich explants made from embryos at time points, immediately after construction, assuming no change upon explantation, for purposes of comparison (B). Giant sandwich explants were made prior to the onset of gastrulation, allowed to heal under a coverglass for 1 hour, then allowed to equilibrate for at least 30 minutes after coverglass removal, prior to fixation. Explants were then cut parasagittally in the dorsal and ventral regions and the resulting “chops” placed on edge and imaged via confocal. Measurements of the total thickness of the lower and upper IMZ and NIMZ as in the examples shown (C, D), to generate the data shown in Figure 4A. The thickness of superficial epithelial cells overlying the IMZ in such explants, as well as whole embryos, was also measured (not shown); resulting data shown in Figure 7H.

**Figure 8-supplementary figure 1.**
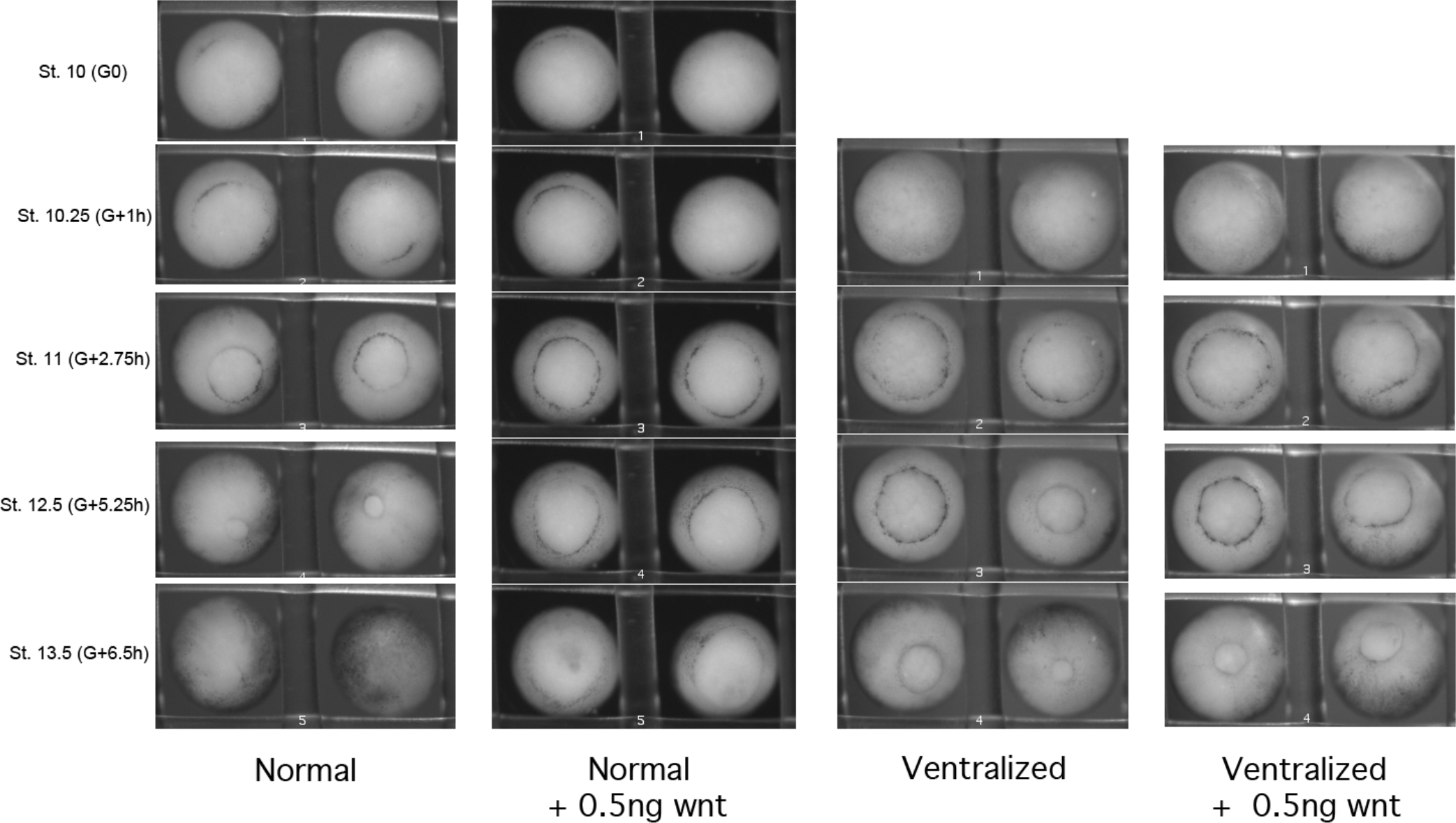

## MOVIES

Movie A. Blastopore closure in normal vs. ventralized embryos. Top two embryos are normal, bottom two were UV ventralized. The two on the left are those shown in Figure 2A. Movie starts at about stage 10 and continues through neurulation (most of which is occurring on the opposite side of the normal embryos).

Movie B. Blastopore closure in normal and ventralized embryos, either uninjected or injected to 10 μMolar MHC II B MO. From left to right: Normal; Ventralized; MHCIIB MO injected; Ventralized & MHCIIB MO injected. Movie starts at about stage 10.5 and continue through neurulation.

Movie C. Blastopore closure in normal and ventralized embryos, either uninjected or injected with 0.5 ng dnWnt.

Movie D. Early normal giant sandwich explant.

Movie E. Early UV ventralized giant.

Movie F. Normal giants with & without bottle cells. Giant with bottle cells on left, without on right. Movies begins about stage 10.25, just after release from coverglass. Time stamp on right movie is fast by 1:02

Movie G. Lithium Dorsoanteriorized giant.

Movie H. Ventral 180° sandwich explant.

Movie I. Dorsal 180° sandwich explant.

Movie J. Double NIMZ and double IMZ sandwich explants.

Movie K. Aggregates of deep cells rounding up.

Movie L. Wontons rounding up.

Movie M. Double IMZ and NIMZ explants from the side.

Movie N. D120 bottle cell retraction and spreading. Nice example of adjacent epithelium being pulled toward forming bottle cells. Starts stage 9+ (G-1.75h) through early neurulation (G+7.75h).

Movie Q1. Keller explant from stage 9.5 to 10+

Movie Q2. Keller explant from stage 10+ to 11

Movie Q3. Keller explant from stage 11 to 12

Movie Q4. Keller explant from stage 9.5 to XX

Movie R. Epithelial spreading/contraction assay. Figure 6A

Movie S. Epithelial retreat and respreading. Movie from which Figure 6C was taken. Animal cap epithelium, recombined with ventral deep IMZ and NIMZ; vegetal end on left. Under coverglass the entire time. Starts stage 10.25 through early neurulation. 3 minutes per frame, 150 frames, 15 fps.

Movie T. High resolution movie of VMZ engulfment & cell motility. Both larger, initially more vegetal IMZ cells and smaller animal IMZ cells show motility over NIMZ region. Vegetal toward the bottom A small patch of epithelium (cells much less motile) appears on the IMZ on the right side about 1/3 of the way through the movie, is pushed onto the NIMZ, then is eventually covered by IMZ cells. Imaged surface is under agarose, under coverglass. Made with a 10x dipping lens on an Olympus AX70. Starts stage 10.25 through early neurulation. 1.5 minutes per frame, 250 frames, 15 fps.

Movie U. Naked dorsal IMZ showing engulfment and reversal. IMZ cells begin to engulf NIMZ, then, at stage 10.5 (15:00), stop and reverse course, eventually converging and extending. Under coverglass; imaged at 10X on an Olympus IX70. Starts stage 10.25 through early neurulation. 1.5 minutes per frame, 250 frames, 15 fps.

Movie V. Naked ventral IMZ showing engulfment. IMZ cells engulf NIMZ; no reversal at stage 10.5 (15:00). Under coverglass; imaged at 10X on an Olympus IX70. Starts stage 10.25 through early neurulation. 1.5 minutes per frame, 250 frames, 15 fps.

Movie W. Ventral giant sandwich. Bottle cells not included. Begins stage 9+ (G-1.5h) through early neurulation (G+7.45). 3 minutes per frame, 180 frames, 15 fps.

